# Transcription termination by RNA polymerase I

**DOI:** 10.1101/2023.11.24.568579

**Authors:** Tomasz W. Turowski, Elisabeth Petfalski, Marie-Luise Winz, David Tollervey

## Abstract

Transcription elongation is stochastic and driven by a Brownian ratchet mechanism, making it subject to changes in velocity. However, on regions occupied by multiple polymerases, notably the rDNA, DNA rotation plus torsion constrain polymerase molecules to proceed at the same rate generating “torsional entrainment”. We report that release of entrainment, by co-transcriptional 3’-end cleavage, is permissive for relative movement between polymerases, promoting pausing and backtracking. Subsequent termination (polymerase release) is facilitated by the 5’-exonuclease Rat1 (Xrn2) and backtracked transcript cleavage by RNAPI subunit Rpa12. These activities were reproduced *in vitro*. Short nascent transcripts close to the transcriptional start site, combined with nascent transcript folding energy, similarly facilitate RNAPI pausing. Nascent, backtracked transcripts at pause sites, are targeted by both the exosome cofactor TRAMP and Rat1, promoting termination. Topoisomerase 2 localizes adjacent to RNAPI pause sites, potentially allowing continued elongation by downstream polymerases. Biophysical modeling supported substantial (∼10%) premature termination.

**Highlights:** Nascent pre-rRNA 3’ cleavage promotes RNAPI deceleration and termination RNAPI undergoes early, start-site proximal termination at sites of polymerase pausing Biophysical modeling indicates ∼10% early termination – or ∼100 events per minute Model presented for overall organization of pre-rRNA transcription

## INTRODUCTION

The pre-ribosomal RNA (pre-rRNA; 7kb in yeast, 10kb in humans) encodes the 18S, 5.8S and 25S/28S rRNAs flanked by the 5’ and 3’ external transcribed spacers (5’ETS and 3’ETS) and separated by internal transcribed spacers 1 and 2 (ITS1) and ITS2) (Fig. 1A and Fig. S1). Ribosome synthesis initiates co-transcriptionally, with assembly of the nascent pre-ribosomes during pre-rRNA transcription (Fig. 1B). In actively growing yeast, each transcribed ribosomal DNA (rDNA) is typically associated with around 50 RNA polymerase I (RNAPI) molecules (Fig. 1B). Transcription elongation is composed of many successive cycles of nucleotide addition, in which the translocation step is based on Brownian motion without input of external energy. Dependence on this “Brownian ratchet”, rather than an energy-driven processive mechanism, makes elongation prone to frequent backtracking and potentially sensitive to inhibition or acceleration by quite modest forces ^1–3^. In the case of RNAPII, pausing and backtracking is common and a major factor in overall gene expression (see for example: ^4,5^.

**Figure 1.**
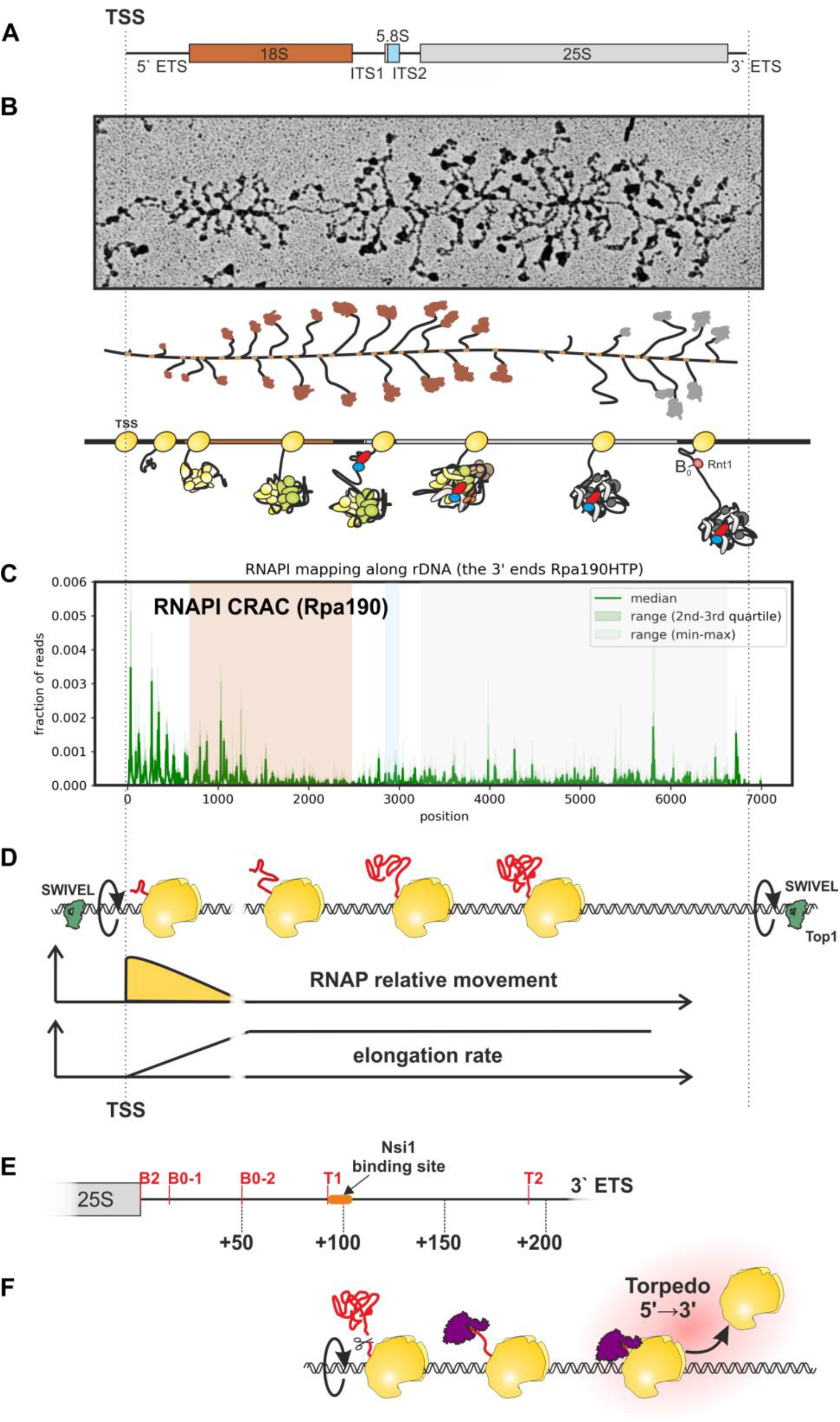
RNAPI convoys are torsionally entrained on spinning DNA. A: Schematic of the 35S pre-rRNA. B: Image and schematics of the rDNA with RNAPI and nascent pre-ribosomes. C: Distribution of RNAPI CRAC reads along the rDNA, highlighting the peak at the 3’ end of the 35S pre-rRNA. Rpa190-HTP CRAC data re-processed from ^7^. D: Cartoon indicating the modeled RNAP elongation rate along the rDNA. Top panel: RNAP are torsionally entrained on the DNA, which is spinning rapidly (∼230 rpm) during transcription elongation. Topoisomerase I (Top1) sites flanking the transcription unit release DNA torsion, introducing single strand nicks in DNA ^30–32^. Middle panel: Cartoon demonstrating implementation of Low Entrainment Region (LER): Initial ability of RNAP to spin at the beginning of the transcription unit. Bottom panel: Consequence of LER is progressive increase of RNAP velocity at the beginning of the transcription unit. E: Scheme showing processing sites within the 3’ ETS: B2 site at the very 3’ end of 25S rRNA, two B0 sites that are cleaved together across a stem structure by the RNase III homologue Rnt1, termination sites T1 and T2, and binding site of road-block terminator Nsi1. F: Cartoon illustrating Rat1-Rai1 torpedo termination of RNAPI transcription.

We previously determined the distribution of transcribing RNAPI across the rDNA, comparing the densities revealed by analyses of “Miller” chromatin spreads, RNAPI chromatin immunoprecipitation (ChIP) and the CRAC crosslinking technique ^6,7^. Two key features emerged; there was an apparent excess of RNAPI over the 5’ region of the pre-rRNA and a very uneven polymerase distribution, with marked throughs and peaks particularly over the 5’ region (Fig. 1C). The strikingly uneven distribution of transcribing RNAPI (Fig. 1C) largely reflected positive effects of nascent RNA folding in promoting rapid transcription elongation, with peaks of polymerase density (slow elongation) correlating with weak folding in a 65 nt window of extruded RNA behind the polymerase (^7^; reviewed in ^8^). This was strongly modulated by the effects of torsion generated by the requirement for one complete rotation of the DNA (or of the polymerase plus nascent pre-ribosome around the DNA) for each ∼10.3 nt synthesized. The megadalton-sized transcribing RNAPI-pre-ribosome complexes are generally very much larger than the DNA separating adjacent polymerases, so the array of polymerases will generally spin the rDNA. At the reported elongation (∼40 nt sec^−1^) ^9^, this will occur at around 230 rpm, with ∼680 complete rotations required for each pre-rRNA molecule transcribed. Since the large, nascent pre-ribosomes effectively block rotation by the polymerase complex around the DNA, the relative movement of the many polymerases on the same rDNA is locked by the buildup of torsion between each of them. This “torsional entrainment” has less effect over the 5’ region of the rDNA, since the short nascent transcripts allow greater freedom for rotation by the polymerases, permitting relative movement along the rDNA (Fig. 1D) ^7^. This is permissive for changes in the relative positions of the polymerase, allowing the large peaks and troughs in RNAPI density seen over the initiation proximal region (Fig. 1C). Consistent with strong torsion, RNAPI transcription also requires topoisomerase activity, with elongation blocked in the 5’ region of the rDNA by combined depletion of Top1 and Top2 ^6,10^. Top1 binds the rDNA both 5’ and 3’ to the transcription unit, recruited by the DNA binding protein Fob1 ^11–13^. We postulate that Top1 at these sites acts as swivels, allowing the entire rDNA repeat to spin under the combined effects of the many engaged RNAPI complexes (Fig. 1D).

Previous analyses of yeast RNAPI transcription *in vitro* and *in vivo* in yeast showed that ∼90% of transcripts terminate at site T1, a T-rich element located at ∼93 nt downstream from the 3′ end of the 25S sequence (site B2) (Fig. 1E) ^14–17^. Termination requires co-transcriptional cleavage by Rnt1, an RNase III family member, across a stem–loop structure within the 3′-ETS, at positions +14/15 and +49/50 (sites B0-1 and B0-2) relative to the 3′ end of the 25S rRNA sequence (Fig. 1E) ^18–20^. B0 cleavage allows entry of the processive nuclear 5′–3′ exonuclease Rat1 (Xrn2 in humans), which degrades the nascent transcript and provokes “torpedo” termination (Fig. 1F). Binding by <u>N</u>TS1 <u>si</u>lencing protein 1 (Nsi1) at a site 12–20 nt downstream from T1 also contributes to efficient termination ^14,21–23^.

Deletion of the *RPA12* gene, which encodes a small subunit of RNAPI, also leads to increased read-through of T1 ^15^. Rpa12 is implicated in RNAPI transcription initiation, elongation and in the cleavage of the nascent transcript when this is in a backtracked position ^24–26^. Deletion of only the C-terminal domain of Rpa12 separates these functions, with a loss of cleavage activity but minimal effects on RNAPI core activity ^27–29^.

Here we report that 3’ cleavage of the nascent transcript, which should release torsional entrainment by allowing polymerase rotation, leads to slowed elongation and favors termination by creating an entry site for the Rat1 exonuclease. Moreover, analyses of RNA surveillance factors and adenylated nascent transcripts indicates that polymerase pausing early in the transcription unit is associated with termination. Stalling of a single RNAPI molecule can potentially disrupt transcription of an entire rDNA unit. However, CRAC using Top2 showed binding to 5’ ETS transcripts, adjacent to stall sites, suggesting a role in breaking torsional entrainment. A biophysical model indicates that RNAPI stalling occurs on around 10% of pre-rRNA transcripts. The conclusion that transcription stalling is relatively common has implications for the overall organization of pre-rRNA synthesis.

## RESULTS

### RNAPI distribution indicates very rapid transcription termination following 3’ cleavage

During pre-rRNA transcription, multiple polymerases on each active rDNA from a “convoy” moving together. Changes in the elongation rate of a single RNAPI molecule within the convoy requires it to rotate relative to the rest of the convoy. Over much of the rDNA, RNAPI is linked to very large pre-ribosomal complexes, greatly constraining such rotation. However, short nascent transcripts present on RNAPI in the 5’ ETS shortly after transcription initiation, allows greater freedom for rotation around the rDNA and relative movement (Fig. 1D) ^7^. This allows greater variability in elongation rates within this region (shown by heterogeneous RNAPI occupancy), which is coupled to folding of the nascent transcripts (Figs. 1C).

RNAPI can be mapped with nucleotide resolution at the 3’ ends of nascent transcripts, using UV-crosslinking and analysis of cDNAs (CRAC) in strains expressing Rpa190 (catalytic subunit) or Rpa135 as fusions with a C-terminal His6-TEV-protein A (HTP) affinity purification tag ^7^. This revealed a clear peak of polymerase density in the 3’ ETS (green line in Fig 2A). Previous analyses revealed a strong correlation between the rate of transcription elongation and strength of RNA folding within the extruded nascent transcript ^7,8^. This correlation was particularly strong over the 5’ ETS region but was clearly visible across the pre-rRNA. The 3’ ETS region was a marked exception, since there is strong folding of the nascent transcript (low ΔG; yellow line in Fig. 2A) flanking a site of high RNAPI occupancy and therefore low elongation rate (green line in Fig 2A). A stable stem is certainly present at this position *in vitro*, since it forms the extended, double-stranded structure essential for Rnt1 cleavage ^18–20^. Note that the yellow line shows the folding energy for the 65 nt of extruded RNA located 5’ to each position – not the folding energy at the site.

**Figure 2.**
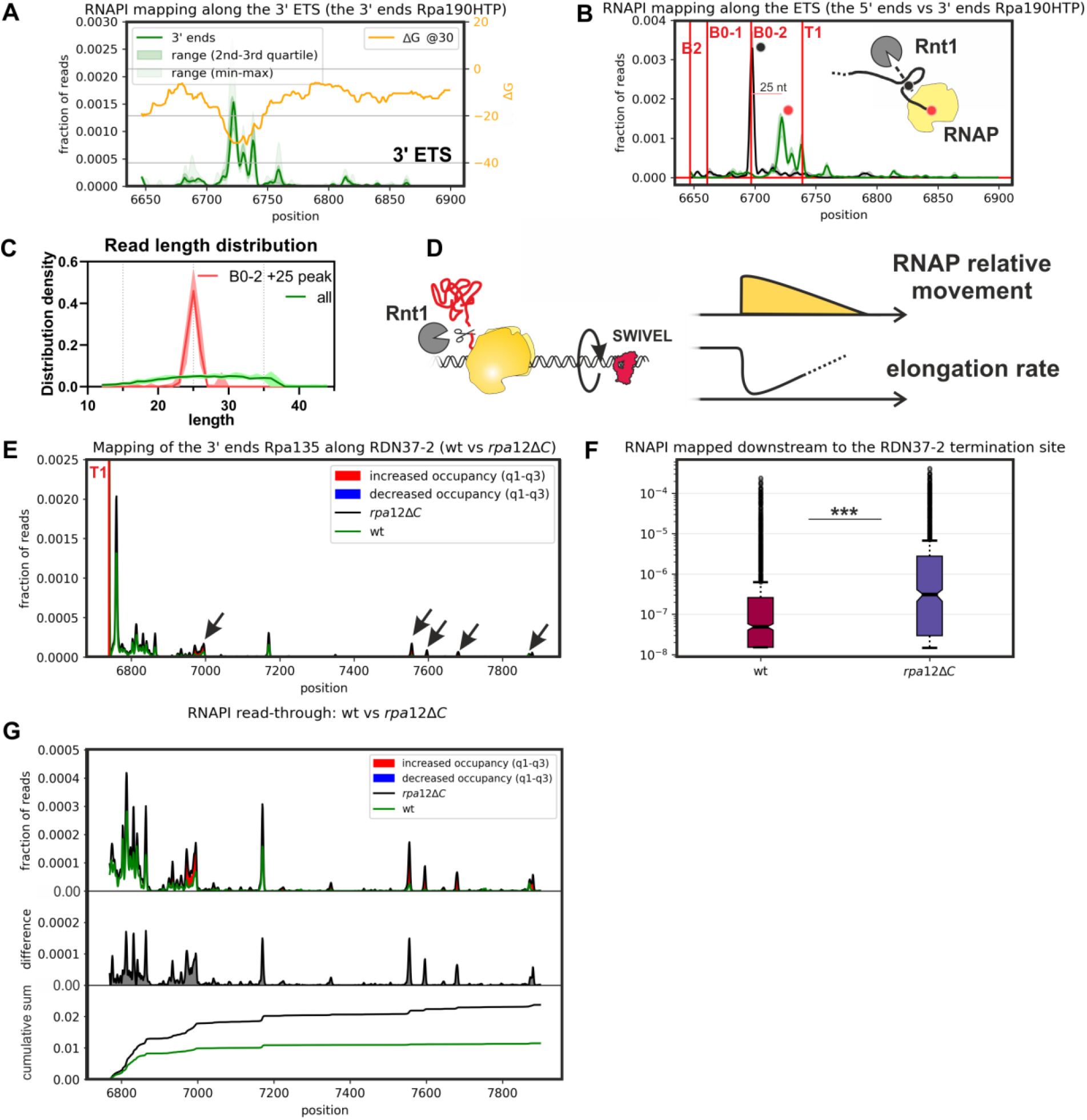
RNAPI termination is associated with decreased elongation kinetics. A: Green line; distribution of RNAPI CRAC reads around the 3’ end of the 35S pre-rRNA. Yellow line; folding energy in a rolling window of 65 nt, offset by 15 nt, behind the polymerase at each nucleotide position. B: Distribution of RNAPI CRAC reads around the 3’ end of the 35S pre-rRNA. Rnt1 processing sites (B0-1, B0-2) and the reported major terminator (T1) are indicted. Black line; distribution of 5’ ends of RNAPI CRAC reads around the 3’ end of the 35S pre-rRNA. The major peak corresponds precisely to the 3’ B0 cleavage site. Green line; distribution of RNAPI CRAC reads 3’ ends. The peaks of the 5’ and 3’ ends distributions are positioned 25 nt part. Cartoon showing the two B0 sites that are cleaved together across a stem structure by the RNase III homologue Rnt1. Peaks corresponding to the major 5’ (black) and 3’ (green) peaks were labelled on the plot and the cartoon with round markers. C: Read length distribution plot for all reads mapped to *RDN37-2* (all) and reads mapped at the major peak within the 3’ ETS (B0-2 +25 peak), marked with green on the panel B. D: Cartoon indicating decreased RNAPI elongation rate as a result of Rnt1 cleavage. RNAPI is released from torsional entrainment on the DNA by the Rnt1 cleavage (top panel). This facilitates rotation of RNAPI together with the DNA (middle panel) and allows a decreased elongation rate (bottom panel). E: RNAPI (Rpa135-HTP) CRAC shows increased readthrough in a *rpa12ΔC* strain, in which endonuclease cleavage of backtracked transcripts is abrogated. F: Boxplot showing fraction of reads downstream to the RDN37-2 termination site for wt and *rpa12ΔC* strains (p-value <0.001, two-sided Wilcoxon test).

Within the 3’ ETS, the peak of polymerase density is located between the B0 cleavage site and terminator T1. To better understand this location, we mapped the 5’ ends of CRAC reads (black line in Fig. 2B and Fig. S2A-B). This revealed a very strong peak at the 3’ Rnt1 cleavage site (B0-2, black dot). Strikingly, reads with 5’ ends at site B0-2 had a median length of 25 nt (horizontal line in Fig. 2B). This would be consistent with very rapid co-transcriptional cleavage of the nascent pre-rRNA by Rnt1 (Fig. 2C). We cannot formally exclude the possibility that some Rnt1 cleavage occurs post-lysis, but the low temperature (ice or 4°C) and absence of Mg^2+^ in the lysis buffer makes this less likely. Notably, the major RNAPI associated peak of B0-2 cleaved RNA did not extend to the reported termination site at T1 (red line in Fig. 2B), indicating that the 3’ end is a site of transcriptional pausing or backtracking. Nascent transcript cleavage was previously reported at site T1 ^33^, but this was not evident in our data as no clear 5’ end peak was observed at this position.

Transcription termination has previously been linked to pausing and backtracking of the polymerase ^34^. This is suppressed by torsional entrainment, since each polymerase complex is “pushed” and “pulled” by all others on the transcription unit (typically around 50 for the rDNA) ^7,35–37^. In the 3’ ETS region the pre-rRNA undergoes co-transcriptional cleavage by Rnt1. We postulate that release from torsional entrainment, associated with the resulting very short nascent transcripts, would facilitate transcriptional pausing/backtracking, and might contribute to termination (Fig. 2D).

Deletion of the gene encoding Rpa12 was previously shown to increase transcription readthrough of site T1 in Miller spreads ^15^ but also greatly reduces overall pre-rRNA synthesis. Rpa12ΔC, deleted for the C-terminal domain, lacks nascent transcript cleavage activity but retains Rpa12 function in RNAPI elongation ^27–29^. We therefore assessed the effects of Rpa12ΔC truncation on RNAPI occupancy across the rDNA. In these analyses, HTP-tagged Rpa135 ^7^ was used to map RNAPI (Fig. S2C and S2E). Over the 3’ ETS, strains with *rpa12ΔC* showed an elevated CRAC signal for RNAPI across intergenic spacer region 1 (IGS1) 3’ and site T1 (Fig. 2E and 2G; quantified in Fig. 2F and Fig. S2F). We conclude that cleavage of the backtracked nascent transcript within RNAPI contributes to efficient termination. However, the major RNAPI peak of occupancy remained downstream of the Rnt1 cleavage sites.

The DNA-binding protein Nsi1 binds a region 12–20 nt 3’ to site T1 ^14,21–23^ and mediates “road-block” termination. To assess the extent to which the 3’ ETS peaks of RNAPI occupancy reflect pausing enforced by Nsi1, RNAPI CRAC was repeated in a *nsi1Δ* strain and in a *nsi1Δ, rpa12ΔC* double mutant (Fig. S2). Relative to the wild-type, RNAPI occupancy downstream of site T1 was elevated in the *nsi1Δ* strain, and further increased in the double mutant (Fig. S2G-I). However, the strong peak of RNAPI occupancy upstream of site T1 remained in all strains.

We conclude that both nascent transcript cleavage by Rpa12 and transcription pausing promoted by Nsi1 contribute to efficient termination, in addition to degradation by Rat1. The peak of RNAPI density downstream of the Rnt1 co-transcriptional cleavage site was not strongly dependent on the presence of Nsi1, consistent with a large contribution from loss of torsional entrainment following pre-ribosome release.

### Reconstitution of transcription termination *in vitro*

To better characterize the role of nascent RNA degradation in triggering RNAPI termination, we developed *in vitro* system (outlined in Fig. 3A and shown in more detail in Fig. S3). We previously reported a system for *in vitro* transcription and backtracking by RNAPI, that was analyzed by following the RNA products of the reactions ^7^. We designed a strong structural element in the nascent transcript which allows to block backtracking of the polymerase and subsequently exchange reaction buffer. To assess transcription termination, we modified this system to measure release of the polymerase from the DNA template, in addition to the RNA product. To achieve this, we immobilized the double stranded template DNA on streptavidin beads *via* template strand, allowing termination to be monitored by release of the transcribing polymerase into the supernatant. The elongating transcription complex was stalled and allowed to backtrack, followed by degradation of the nascent transcript. The 5’ exonuclease Rat1 normally functions in a stoichiometric complex with the pyrophosphatase Rai1 ^38,39^, and we therefore purified the Rat1-Rai1 complex via TAP-tagged Rat1. For *in vitro* transcription we generated a construct with a G9 element (Fig. S3A), and induced stalling by transcription in the absence of GTP. Rat1 is most active on substrates with a 5’-monophosphate, so this was added to the transcription primer by polynucleotide kinase (PNK) treatment (Figs. 3A and S3B).

**Figure 3.**
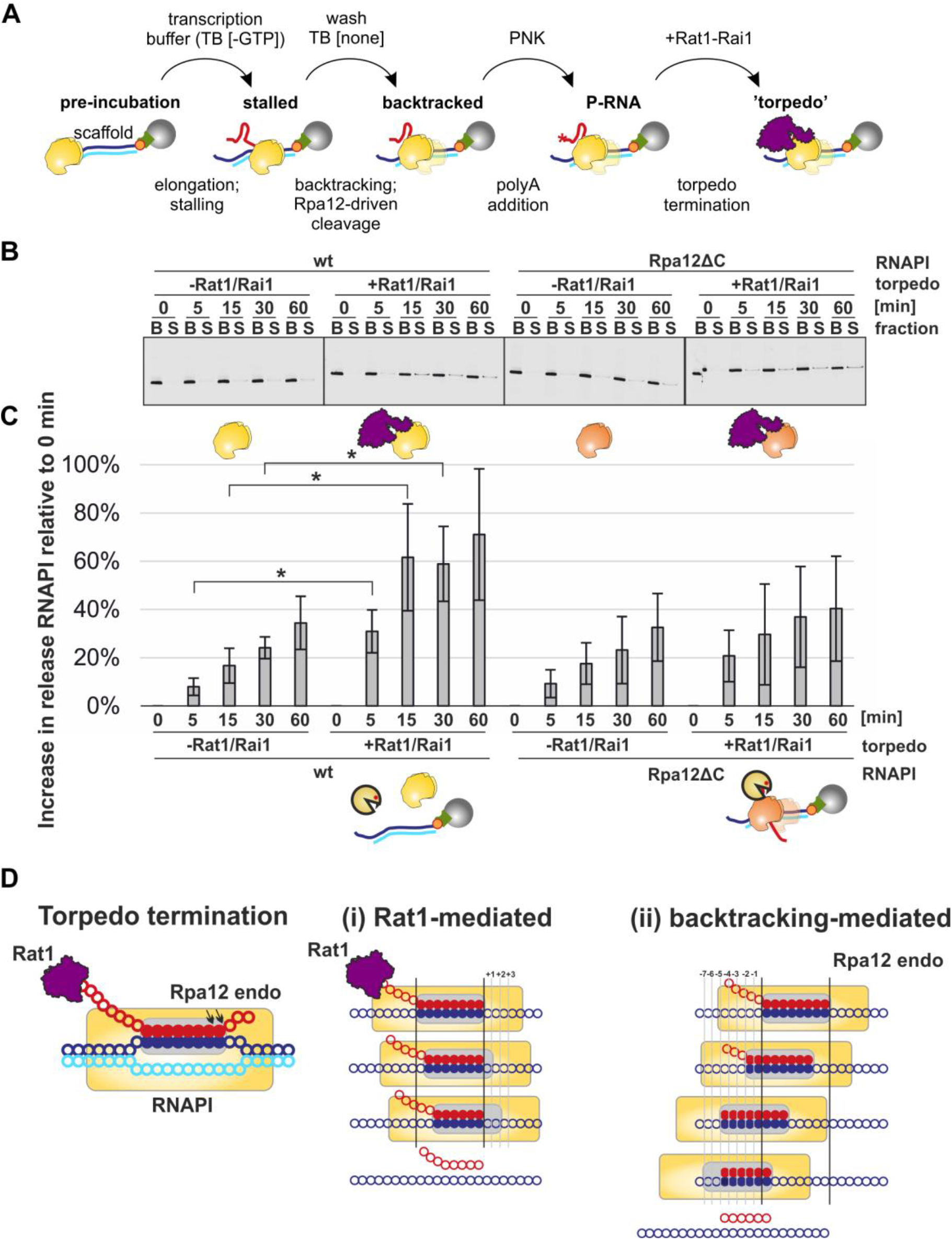
Efficient termination requires exonuclease activity by Rat1 and endonuclease activity of RNAPI. A: Schematic of *in vitro* termination assay. Description in text. For detailed version see Fig. S3. B: *In vitro* termination assay. Western blot showing Rpa135-HTP (RNAPI) distribution between the DNA-bound (B) fraction and the supernatant (S) that represents terminated and released RNAPI. C: Quantitation of data in panel B. Some release of RNAPI from the template DNA was already observed at T0, so this initial level was set to 0. The subsequent increase in RNAPI release over time is indicated by the graphs. See methods for the details. Asterix indicates significant differences (p-value < 0.05, two-tailed T-test, n=5). D: Two potential mechanisms participating in torpedo termination of RNAPI: (i) Rat1-mediated mechanism and (ii) backtracking-mediated mechanism.

Recovery of RNAPI in the streptavidin/DNA bound fraction (B; not terminated) and supernatant fraction (S; terminated) was compared during a time course of Rat1/Rai1 treatment (Fig. 3B; quantified in Fig. 3C). This showed a significant increase in termination induced by transcript degradation. Rpa12 is an integral component of RNAPI. To assess the role of nascent transcript termination *in vitro*, the assay was repeated using RNAPI purified from a *rpa12ΔC* strain. Using the Rpa12ΔC form of the polymerase, significant enhancement of termination was not seen following Rat1/Rai1 treatment (Figs. 3B and 3C), consistent with our *in vivo* data (Figs. 2E and 2F).

These results show that both Rat1 and Rpa12 activities are needed for efficient torpedo termination *in vitro* (Fig. 3D). Rat1 shows processive 5’-exonuclease activity *in vitro* ^40^. In principle, the RNAPI elongation complex could be “pushed” forwards, without nucleotide incorporation, to the position at which the transcription bubble becomes unstable and favors RNAPI dissociation (Fig. 3D, panel (i)). However, this requires DNA strand separation without compensatory RNA-DNA base pairing, and it is unclear whether sufficient force can be generated by Rat1-Rai1. We propose an additional mechanism, in which the torpedo trims nascent RNA only to the boundary of the elongation complex, with further steps based on a backtracking-mediated mechanism until transcription bubble becomes unstable (Fig. 3D, panel (ii)). This mechanism would be favored by the elongation complex which can ‘slide’ backwards on the DNA. There is, however, competition between transcript elongation and endonuclease activity of the polymerase.

### Early termination by RNAPI

Termination in the 3’ ETS, the end of the transcription unit, appears to be associated with polymerase pausing and/or backtracking. Previous studies also indicated substantial peaks of RNAPI occupancy in the 5’ ETS, which we interpreted as indicating pause sites ^6,7,41^. Our previous analyses indicated that the heterogeneity in elongation rates was linked to the reduced levels of torsional entrainment allowing the effects of nascent RNA folding to be strongly manifested. However, they did not exclude the possibility that some premature termination also occurs for RNAPI, as previously reported for RNAPII ^42,43^.

Premature transcription termination is expected to be associated with degradation of the nascent transcript. During normal pre-rRNA processing, the excised 5’-A0 region of the 5’ ETS is degraded by the exosome together with the DExH RNA helicase Mtr4 ^44,45^, which is directly recruited by the pre-ribosome component Utp18 ^46^. RNA surveillance activities of the yeast nuclear exosome require the TRAMP (Trf4/5-Air1/2-Mtr4 polyadenylation) complex, which facilitates RNA degradation by addition of a single stranded oligo(A) tail. However, TRAMP components other than Mtr4 are not required for pre-rRNA processing (5.8S trimming and 5’ ETS degradation). The 5’ exonuclease Rat1 is not a major factor for normal degradation of the 5’-A0 region, probably reflecting a degree of protection conferred by the 5’ tri-phosphate on the primary transcript ^47^, but degrades the excised A0-A1 region ^48^.

Mapping the distribution of Rat1 across the 5’ ETS revealed a pattern of peaks (Fig. 4A and Fig. S4A). Reads were mapped using the 5’ ends of the recovered sequences, since this expected to correspond to the location of Rat1. The strongest peak was at +610, corresponding to the 5’ end of the excised A0-A1 pre-rRNA region. Since yeast rDNA sequences are generally identical, metagene analyses cannot be applied. However, we have previously used a peak-calling algorithm to identify common features at different sites within the rDNA ^7^(outlined in Fig. S4B). Using this approach, we identified and aligned peaks (except peak 643, Fig.S4B) of occupancy for Rat1 (Fig. 4B; purple line) and RNAPI (Fig. 4B; green line). Rat1 occupancy showed a peak ∼40 nt upstream from the peak of RNAPI occupancy, mapped using the 3’ ends of the sequences. We interpret this as showing Rat1-Rai1 degradation of nascent transcripts associated with stalled or paused RNAPI – consistent with the occurrence of torpedo termination within the 5’ETS.

**Figure 4.**
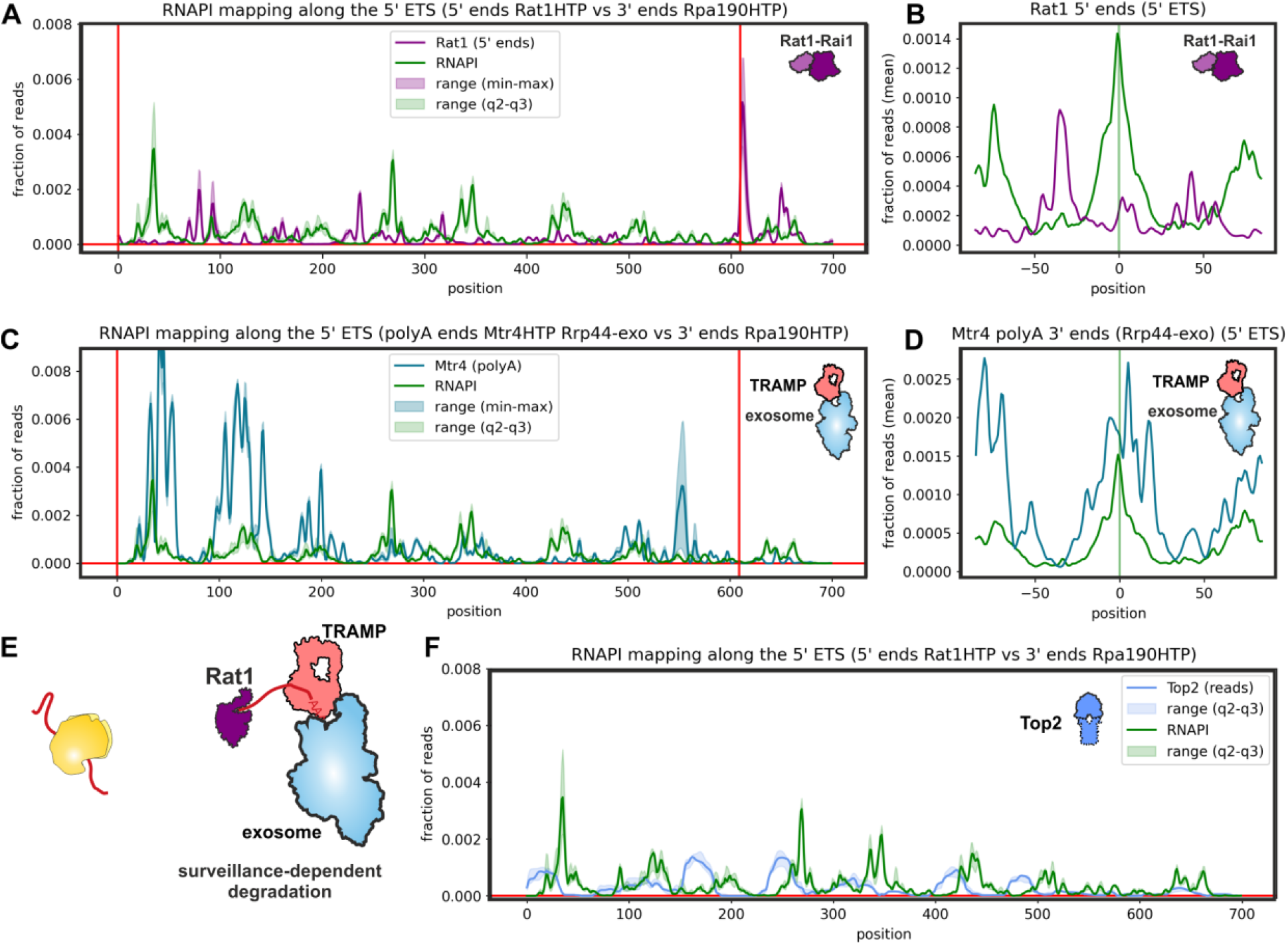
Rat1 and TRAMP are found at sites of slowed RNAPI elongation. A: Rat1 found at sites of slowed / paused RNAPI. The 5’ ends of mapped reads are presented. B: RNAPI CRAC peak metaplot for RDN37, comparing Rpa190 and Rat1 at the level of single peaks. Note: +643 peak was excluded from this analysis (see also Fig. S4B). C: TRAMP found at sites of slowed / paused RNAPI. The figure shows 3’ ends of mapped, oligo(A) reads recovered in an Rrp44-exo strain background. D: RNAPI CRAC peak metaplot for RDN37, comparing Rpa190 and the 3’ oligo(A) ends of Mtr4 reads, at the level of single peaks. Note: +643 peak was excluded from this analysis. E: Cartoon proposing mechanism how surveillance-liable fragments could be generated. Prolonged pausing of RNAPI would induce Rpa12-mediated cleavage and release of ∼40 nt-long RNA fragments. These would be susceptible to Rat1 and TRAMP targeting. F: Top2-HTP CRAC reads distribution across the 5’ ETS harmonizes with RNAPI pausing/stalling sites.

RNAPI peaks in the 5’ ETS correspond to sites of high occupancy, reflecting greatly slowed or paused elongation. We speculated that backtracking at these sites might expose the 3’ ends of nascent transcript to the surveillance machinery (Fig. 4E). TRAMP-mediated surveillance tags substrates with oligo(A) tails, so we sought these in RNAPI-associated nascent transcripts (Figs. 5A-B, S5A-B). The fraction of RNAPI associated reads that carried non-templated oligo(A) sequences (AAA or longer) was highest at sites of slowest elongation (Fig. 5B). The effects were strongest over the 5’ ETS (Fig. 5A-B), but were visible across the entire rDNA (Fig. S5A-B). We conclude that the nascent transcript can be adenylated while bound to RNAPI, and this is most common at sites where elongation is slowest and prone to backtracking.

**Figure 5.**
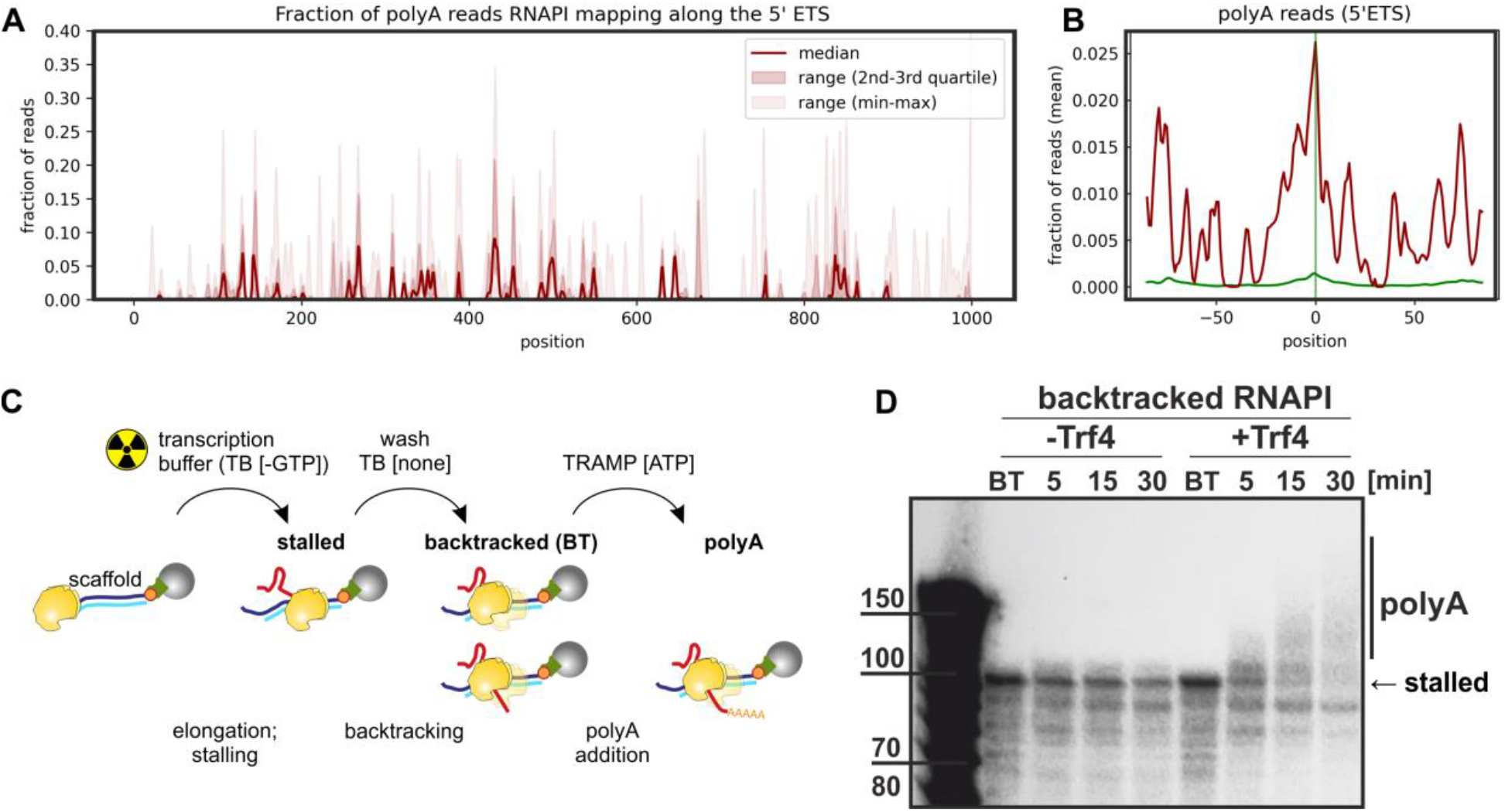
Nascent transcripts can be oligo-adenylated by the TRAMP complex. A: Fraction of oligo(A)+ reads recovered in RNAPI CRAC data (non-coded A_n_ ≥ 3), mapped along the 5’ ETS. B: RNAPI CRAC peak metaplot for 5’ ETS, comparing the 3’ ends of the reads (green) with poly(A) reads. C: Schematic of *in vitro* assay for transcription, backtracking and adenylation. D: Purified Trf4-Air4/5 can add poly(A) tails to nascent RNA associated with backtracked RNAPI. Transcripts were internally radiolabeled and visualized with an Image Analyzer (Fuji-Film).

To assess whether the tails were added by the TRAMP complexes, we mapped Mtr4 binding, mapping the 3’ ends and selecting only oligo-adenylated reads (Figs. 4C and 4D) (^49^ GSE77863). These are expected to reflect sites of TRAMP activity. This analysis was performed in a strain expressing only Rrp44-exo ^50^, lacking the exonuclease activity of Rrp44, to stabilize the very transient intermediates of degradation. RNAPI peaks overlapped with peaks of p(A)^+^ reads recovered with Mtr4 (Fig. 4C), and this was supported by metaprofile analysis (Fig. 4D). Comparison of RNAPI with all Mtr4 reads is shown in Fig. S4C. From these data, we concluded that backtracked, nascent transcripts associated with stalled RNAPI are targets for the TRAMP complex. We also assessed the effects of *rpa12ΔC* on RNAPI density in the 5’ ETS (Fig. S4D). Increased density was seen at some pause sites, indicating that Rpa12 also participates in RNAPI release from pausing. The effects were modest, but the data are normalized so only differences in relative peak heights can be seen.

These data indicate that a significant level of transcriptional pausing by RNAPI occurs within the 5’ETS region. Upstream polymerases will be blocked by the stall. In addition, since transcribing polymerases are expected to be strongly entrained, downstream polymerases may also be torsionally stalled, potentially imposing considerable strain on the DNA. Human Top2A binds RNA, with specificity for 3’ ends carrying 3’-OH groups, as would be the case for backtracked nascent transcripts. ^51^. Yeast Top2 was identified in a screen for RNA-interacting proteins using UV crosslinking followed by density centrifugation (iRAP) ^52,53^. We therefore speculated that Top2 might be recruited to the 3’ end of the nascent pre-rRNA at sites of RNAPI backtracking to relieve torsional stress. A Top2-HTP strain showed no clear growth defects, indicating that the fusion protein is functional, and gave good signals in CRAC, confirming *in vivo* RNA association. The full Top2-CRAC dataset is available from GEO, with accession number GSE246546. Mapping Top2-HTP crosslinking sites on the rDNA revealed peaks of occupancy in the 5’ ETS that are adjacent to sites of RNAPI pausing/stalling (Fig. 4F). This was supported by metaprofile analyses (Figs. S5E-G). The 3’ ends of oligo(A) tailed, Top2 associated reads mapped more closely with RNAPAI associated oligo(A)^+^ reads (Fig. S5G). This is consistent with Top2 binding to backtracked nascent transcripts.

We propose that Top2 is recruited to stalled, backtracked RNAPI and acts to “break” the torsional entrainment, allowing downstream polymerases to continue transcription.

### Adenylation of nascent transcripts

The overlap between Mtr4 adenylated reads with RNAPI peaks indicated that nascent transcripts can be adenylated while still associated with the transcribing polymerase. In the elongating complex, the 3’ end is sequestered within the polymerase. This suggested adenylation might be specific for stalled and backtracked RNAPI, from which the 3’ end may be extruded as demonstrated for RNAPII ^54^. To test whether TRAMP components can adenylate the nascent transcript associated with backtracked polymerase, we reconstituted this activity *in vitro* (Fig. 5C). The TRAMP4 (Trf4 plus Air1/2) complex was purified from yeast using Trf4-TAP (Open Biosystems). Elongating RNAPI was purified using Rpa135-HTP as above and incubated with the DNA template shown in Fig. S3. RNAPI was stalled at the G9 tract and allowed to backtrack by NTP wash out. Incubation with TRAMP4 plus ATP resulted in progressive elongation of the nascent transcript (Figs. 5D and S5C). When the analysis was repeated using RNAPI-Rpa12ΔC, accumulation backtracked RNA was more pronounced, as expected, but less processive adenylation was observed (Fig. S5C).

The same assay was applied to RNAPII, purified with TAP-tagged Rpb1 (Rpo21). We observed adenylation of the backtracked nascent RNAPII transcript in the presence of purified TRAMP4 (Fig. S5D). For RNAPII, the homolog of Rpa12 is TFIIS (Dst1 in yeast), which is not a stable component of the polymerase holoenzyme. Addition of purified TFIIS reduced backtracked RNA accumulation and adenylation (Fig. S5D).

### *In silico* simulation of premature termination and transcriptional output

The CRAC data strongly suggested that pre-rRNA transcription is subject to a degree of premature termination, but the termination frequency cannot be accurately determined from these data alone. To quantitatively estimate the range of premature termination, we modified a previously developed computational model for RNAPI transcription ^7^. This model incorporates various factors that affect transcription elongation, including stochastic elongation rate, DNA torsion and RNAP entrainment, stability of the transcription bubble within the RNAPI, and the extruded nascent RNA’s impact on RNAP backtracking.

To estimate the range of RNAPI transcriptional processivity consistent with *in vivo* rRNA production, we incorporated literature values, including the presence of 75 transcriptionally active rDNA repeats (50% of the 150 total rDNA repeats) at any given timepoint ^55^, an average of 50 RNAPI per active rDNA repeat ^7,56^, and the production of 200,000 newly synthesized ribosomes per cell, per generation (100 min) ^57^, although this may be considered the higher end of the range ^58^.

We aimed to incorporate different levels of RNAPI processivity into the model and simulated for 6,000 sec (100 min) reflecting one yeast cell division. From the output of 75 repeats, we calculated the total number of fully synthesized pre-rRNA molecules (*RNAP*_productive_), the number of prematurely terminated pre-rRNA (*RNAP*_non-productive_), processivity, and the average number of RNAPI per single rDNA repeat (RNAPI number). The initial results highlighted that premature termination rate cannot be equated directly with processivity, as defined by the ability of RNAPI to reach the end of the transcription unit.

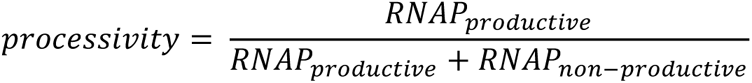

RNAP elongation is a stochastic process that follows a Brownian ratchet mechanism, resulting in a non-uniform distribution of occupancy time, due to multiple factors that affect the velocity. We anticipate that premature termination is related to the occupancy time at a particular position, rather than being equally distributed throughout the transcription unit. This is supported by the observation that low RNAPI velocity (indicating high occupancy) was strongly correlated with RNA transcript binding by the surveillance machinery (Fig. 4).

To account for this, we incorporated the probability of premature termination (*PT*) in our model, which is scaled according to the estimated total transcription time. This approach considers the probability of premature termination at each position along the transcription unit, providing a more accurate representation of the stochastic nature of the transcription process.

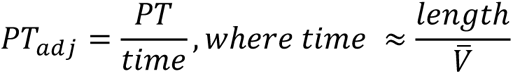

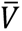 is average velocity. In our implementation, the value of *PT* approximates to *processivity*, although it does not fully satisfy its definition. By incorporating *PT* as a parameter in our model, we consider the likelihood of premature termination at each position along the transcription unit, which provides insight into the stochastic nature of the elongation process. However, *PT* does not directly measure the RNAP ability to reach the end of the transcription unit, which is the definition of *processivity*. Therefore, although *PT* and processivity share some similarities, they are not equivalent measures of RNAP transcription.

We conducted a series of simulations to test the implementation of premature termination as *PT*, using a stochastic RNAP elongation mechanism and an initiation probability of 0.8, as previously. We calculated the number of rRNA molecules produced and RNAPI molecules per rDNA, and investigated how *PT* value influenced RNAPI transcriptional output (Fig. 6A). We previously proposed that the observed 5’ bias in rDNA transcription originates from the progressive entrainment of multiple RNAPI molecules across the transcription unit, with Low Entrainment Regions (LER) close to the transcription start site (TSS) ^7^ (Fig. S6A-C). To validate this model, we included *PT* in the simulations across the first 2,000 nt where 5’ bias was observed (Fig. 6B-C and Fig. S6D-G).

**Figure 6.**
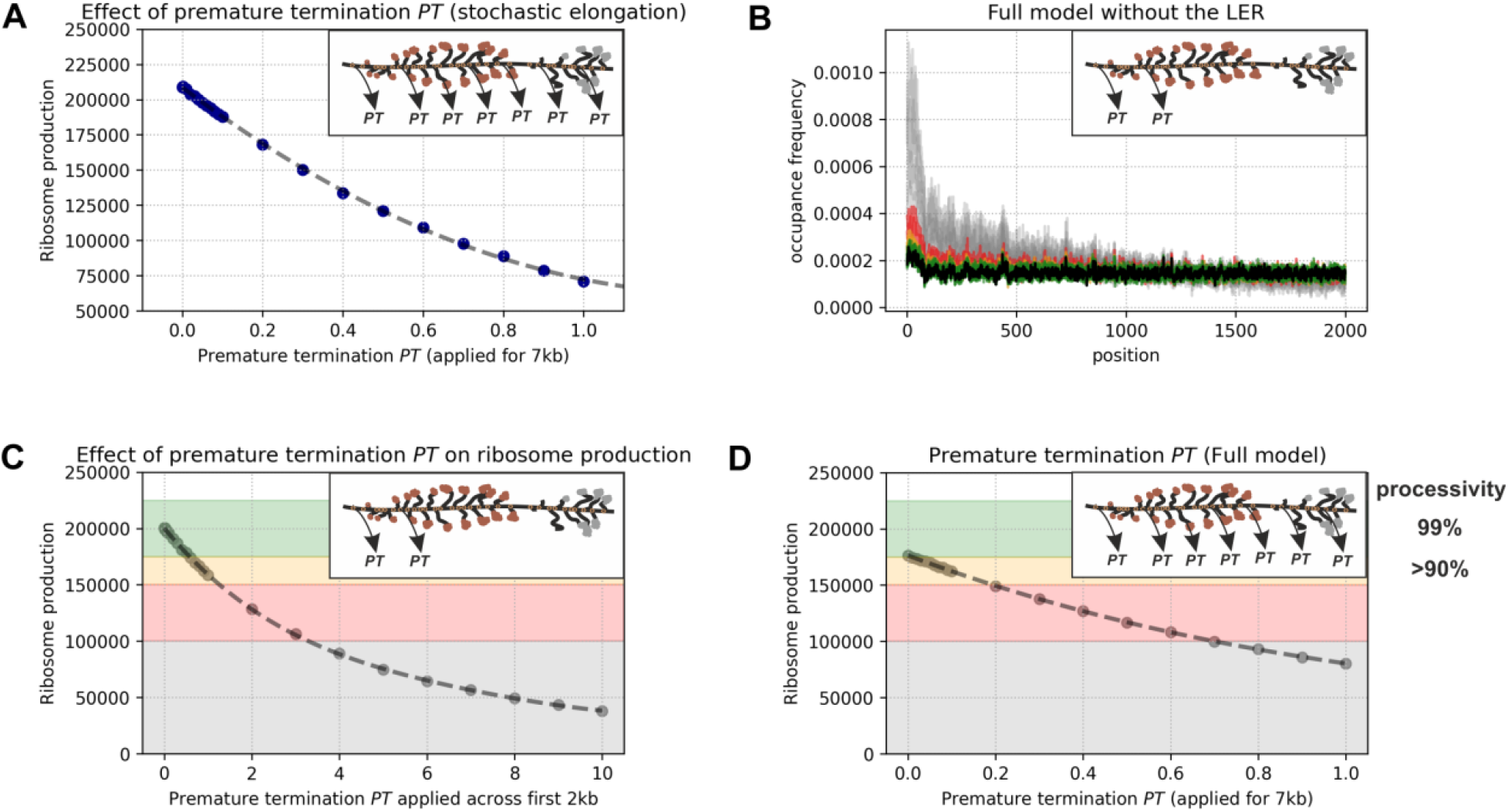
Computational model of RNAPI transcription predicts range of premature termination *in vivo*. A: Effect of premature termination PT on ribosome production in a model where RNAPI velocity is driven by stochastic elongation only. PT was applied across the entire transcription unit. B: RNAPI occupancy profiles after application of PT. Profiles are superimposed and color-coded to correspond with the background on panel C: black – no premature termination PT, green and orange PT where ribosome synthesis is effective, red – ribosome synthesis is decreased (<150k per generation) and grey where ribosome synthesis is insufficient (<100k per generation). C: Effects of PT on ribosome production in a model including stochastic elongation, RNA elements and DNA torsion forces. PT was applied across the initial 2kb of the transcription unit. Background colors match the color-codes on panel B. Note that because PT is applied only across a region of the transcription unit, the values of PT are significantly higher than for panels A and C. D: Effect of premature termination PT on ribosome production. The data indicate that 10% of RNAPI can terminate prematurely without significant impairment of overall ribosome biosynthesis.

We found, that including *PT* in the absence of the LER, was able to reproduce the observed 5’ bias (Fig. 6B), but led to significantly reduced ribosome production (Fig. 6C), which is inconsistent with the demand for 200,000 ribosomes per generation. We concluded that the accumulation of RNAPI signal on the 5’ end of the rDNA transcription unit is unlikely to be solely due to premature termination. We further investigated the range of premature termination that still allows sufficient rRNA synthesis to support cell division (Fig. 6D). PT values below 0.2 were required to produce numbers of ribosomes consistent with the reported range. Whereas values above 0.7 are required to generate the observed excess of polymerases in the 5’ region of the rDNA.

We also observed a significant level of polymerase pausing, which can break the convoy of entrained polymerases and lead to run-off. However, polymerases located 5’ will be blocked and any RNAPI complex that remains stalled will eventually be ubiquitinated and degraded off the DNA ^59,60^. Therefore, the overall frequency of premature termination by RNAPI is unlikely to exceed 20%, but most likely values are around 10%. While this proportion is low, cumulatively 10,000 transcription events may be prematurely terminated per generation (∼100 events per min). At steady state, the cellular abundance of abortive transcripts is likely to be low, due to efficient clearance by the surveillance machinery.

## DISCUSSION

Eukaryotic transcription elongation rates show considerable heterogeneity, with functional consequences for the many RNA processing and packaging factors that act very quickly on the nascent transcript. RNAPI transcribes a single pre-rRNA transcript from the nucleosome-free rDNA, making it well suited to such analyses. We previously observed 5’ enrichment for RNAPI density with strikingly uneven, local polymerase distribution, most notably over the 5’ ETS region of the pre-rRNA (Fig. 1).

In the 3’ ETS the RNAPI associated RNAs showed a strong peak of 5’ ends, precisely at the 3’ cleavage site reported for Rnt1 (B0-2). No clear peak was seen for the 5’ cleavage site, consistent with the expectation that Rnt1, like other RNase III-related enzymes, cleaves across the stem. RNA fragments with 5’ ends at site B0-2, showed a strong bias in length distribution, with 3’ ends greatly favored at position +25 nt. This indicates very rapid co-transcriptional Rnt1 cleavage across the terminal stem, when the polymerase has progressed only 25 nt, with 10-12 nt of extruded nascent transcript. This speed was initially surprising, but we note that pre-mRNA splicing can be comparably rapid *in vivo,* despite much greater complexity. Spliced products are observed as introns emerge from RNAPII and splicing is 50% complete when the polymerase has advanced only 45 nt ^61^. At least in part, this rapidity reflects coupling of slowed/paused RNAPII elongation with splicing ^62,63^. In the case of RNAPI, we postulate that release of the nascent pre-ribosome by Rnt1 cleavage relieves the polymerase from torsional entrainment. This confers increased freedom to rotate with the DNA rather than continuing to elongate. The resulting deceleration may be important in facilitating Rat1-mediated termination. For RNAPII, deceleration following co-transcriptional cleavage plays a major role in making it a “sitting duck” for Rat1 (Xrn2) termination ^64^.

Our data also revealed that endonuclease cleavage by Rpa12 facilitates effective termination, presumably acting on backtracked transcripts. During torpedo termination the nascent transcript has presumably be extracted from the polymerase, and cleavage of backtracked transcripts may facilitate this process. The DNA binding protein Nsi1 was reported to also contribute to termination by binding downstream of the major terminator site T1 ^21–23^. We observed increased readthrough in *nsi1Δ* strains, but the extent was modest and the major peak adjacent to the Rnt1 cleavage site was not clearly altered. Termination may normally be promoted by the interplay between entrainment release following Rnt1 cleavage, roadblocking by Nsi1 and the Rat1 torpedo activity, with Rpa12 cleavage aiding release of backtracked polymerases.

A significant level of transcription start site proximal termination was also identified. We previously reported an excess of RNAPI density, and very uneven RNAPI density within the 5’ ETS ^7^. We concluded that, as for the 3’ ETS, short nascent transcripts in this region facilitate pausing and backtracking, particularly at sites of weak folding in the nascent transcript. At these locations RNAPI is associated with oligo(A) tailed RNAs, indicating 3’ adenylation of backtracked nascent transcripts. Consistent with this the Mtr4 component of the Trf4/5-Air1/2-Mtr4 polyadenylation complex (TRAMP) was localized to transcription pause sites. Moreover, we could reproduce TRAMP-mediated adenylation of nascent transcripts on backtracked RNAPI *in vitro*. These observations all support the model that backtracked, RNAPI is targeted and oligo-adenylated by TRAMP; a key cofactor for the exosome nuclease complex. We predict that the exosome degrades the 3’ extruded pre-rRNAs, potentially promoting termination by a “reverse torpedo” mechanism (Fig. 7). The 5’ exonuclease Rat1 was localized immediately upstream of the RNAPI pause sites, strongly indicating that “conventional” torpedo termination also occurs at these locations. Biophysical modeling is consistent with around 10% of pre-rRNAs transcription terminating in the 5’ ETS. Given the very high rates of ribosome synthesis in yeast, this represents a substantial number of events (around 100 per minute per cell). We cannot accurately determine the relative contributions of forward and reverse torpedo mechanisms to premature termination, but the presumed intermediates in each pathway were detected in unperturbed cells, indicating that both normally participate.

**Figure 7.**
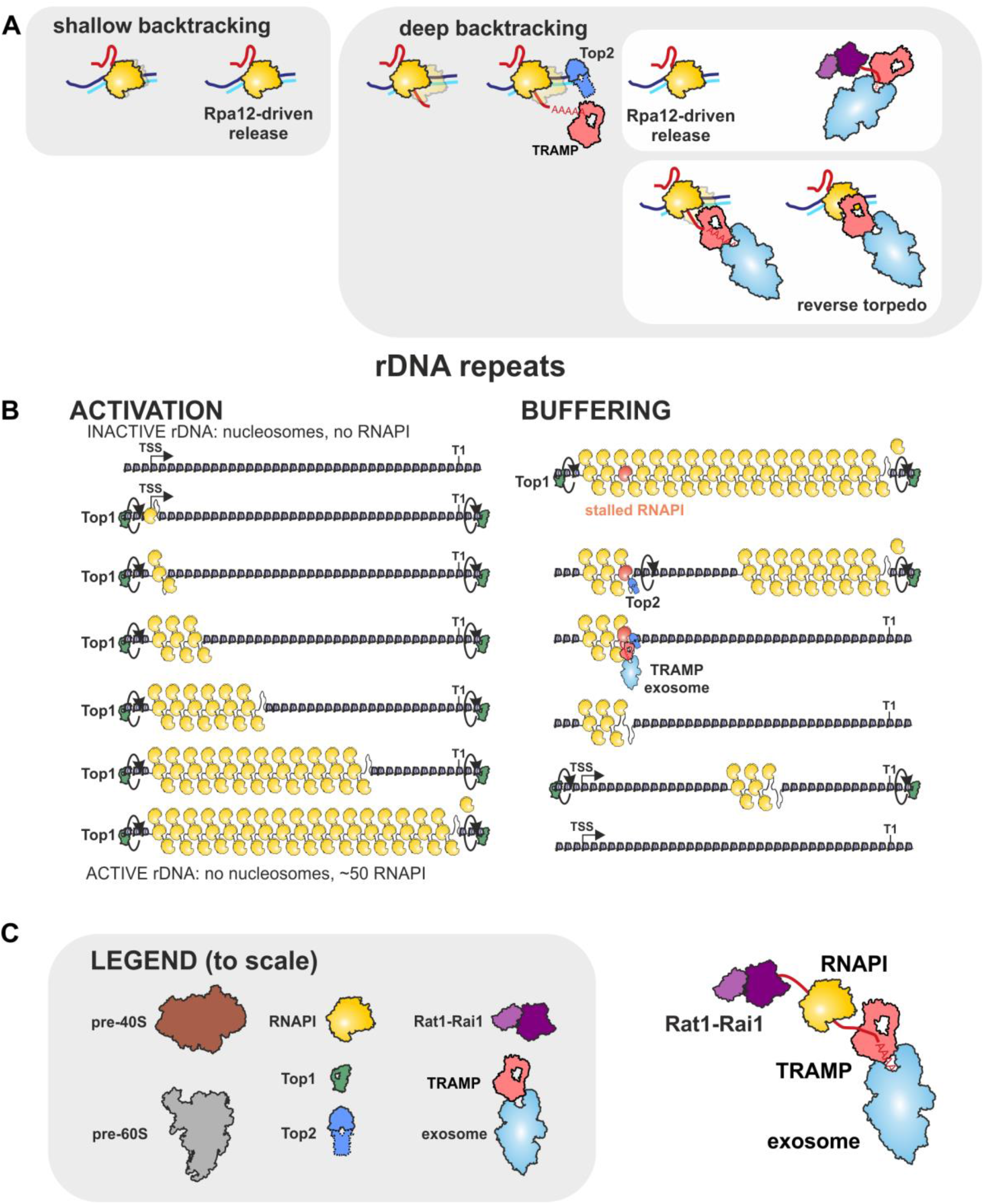
Models for RNAPI transcription A: Shallow backtracking can be released by Rpa12-mediated cleavage of the nascent transcript, but release from deep backtracking is promoted by a reverse torpedo mechanism, initiated by oligo-adenylation by the TRAMP complex. B: Left: Inactive rDNA repeats are packaged in nucleosomes, but can be opened by the cooperative activities of multiple polymerases, each linked by torsional entrainment. Right: A single stalled or deeply backtracked RNAPI will break the convoy of polymerases. Continued transcription by downstream polymerase can be facilitated by Top2 recruitment. The stalled polymerase may be released by combined torpedo (Rat1) and reverse torpedo (TRAMP, exosome) activities. C: Model components summarized to scale, plus the forward and reverse torpedoes.

### Cycle of rDNA transcription

Inactive rDNA repeats are packaged with nucleosomes. In Figure 7 the relative sizes of nucleosomes and RNAPI complexes are approximately correct and consistent with the length of the rDNA. Numbers of molecules per rDNA repeat are also appropriate. The nascent pre-ribosomes would be very much larger in size, with molecular masses up to 10-fold greater than RNAPI - but are omitted for clarity.

In both yeast and human cells, only around half of the rDNA repeats are actively transcribed at any time. The basis for this apparent excess has been unclear, especially in yeast with its compact genome in which the rDNA is around 10% of the total. We postulate that this may be related to the finding of substantial polymerase stalling described here. A single stalled polymerase will break the convoy of coordinately transcribing polymerases on the rDNA, potentially blocking elongation by all polymerases on the transcription unit (Fig. 7).

Shallow RNAPI backtracking (up to 20 nt) can be resolved by the intrinsic endonucleolytic activity of Rpa12 within RNAPI. However, recovery from deep RNAPI backtracking is more limited ^65^ and we propose that this is promoted by the reverse torpedo mechanism (Fig. 7A). If a single polymerase stalls or backtracks, the downstream polymerases, still torsionally entrained, will be moving away at ∼40 nt/sec. So, a second later, around 4 negative supercoils will already have accumulated in front of the stalled polymerase, which can be accommodated by either “writhe” (twisting) or strand separation. We note that the combination of negative supercoils and polymerase backtracking will be conducive to anterior R-loop formation. Consistent with this model, we previously noted a high level of R-loop formation on the rDNA 5’ETS ^66^. With extended pausing, the resulting high torsion will potentially lead to DNA damage. Specific recruitment of Top2 to the backtracked polymerases would allow run off for downstream polymerases, reducing the torsional burden. The stall could then be cleared through termination or, failing that, via ubiquitination and proteosome degradation of the “broken” polymerase ^59,60^.

We postulate that the loss of transcription activity on a single rDNA unit is followed by its closure and packaging into a chromatin array of nucleosomes – accompanied by opening of previously closed repeats to maintain active rDNA repeat numbers. Activation of a repeat requires transcription through the nucleosome array. Initially by a single RNAPI, but as more polymerases are loaded, they will increasingly act cooperatively as a convoy linked by DNA torsion. Transcription by RNAPII through nucleosomes is facilitated by numerous cofactors ^67^, most of which are not shared by RNAPI (but see ^68^). These are required, in part, to re-establish nucleosomes following polymerase passage. However, actively transcribed rDNA regions are believed to be nucleosome free – and the schematic shown in Figure 7 fits with this conceptually. The pioneer RNAPI, aided by all the subsequent polymerases, only needs to displace nucleosomes from the DNA. Nucleosomal DNA is negatively supercoiled, so the positive supercoils expected in front of the leading polymerase may promote nucleosome displacement, facilitating clearance of the transcription unit.

### Data Availability

All sequence data generated for this project are available from the Gene Expression Omnibus (GEO; https://www.ncbi.nlm.nih.gov/geo/), with the reference number GSE246546.

### Author Contributions

TWT and DT conceived the project and wrote the manuscript. EP performed experiments. TWT, EP and DT analyzed the data. TWT and DT developed the mathematical model. All authors edited and reviewed the manuscript.

## Supporting information

Supplementary Informations

## Acknowledgements

We thank Olivier Gadal and Christophe Dez (Centre de Biologie Intégrative (CBI), Toulouse) for critical reading of the MS and Jan Mikołajczyk (IBB PAS) for stimulating discussion during development of the computational model. TWT is supported by the Polish National Agency for Academic Exchange (PPN/PPO/2020/2/00004/U/00001) and National Science Center (2020/39/D/NZ2/02115).

DT and EP were supported by Wellcome Principal Research Fellowships (109916, 222516). Work in the Wellcome Centre for Cell Biology is supported by a Centre Core grant (203149).

## Disclosure declaration

The authors declare that they have no competing interests.

## STAR Methods

### KEY RESOURCES TABLE

| REAGENT or RESOURCE | SOURCE | IDENTIFIER |
| --- | --- | --- |
| Antibodies |  |  |
| TAP Tag Polyclonal Antibody | ThermoFisher | Cat#CAB1001 |
| Bacterial and Virus Strains |  |  |
| Chemicals, Peptides, and Recombinant Proteins |  |  |
| -Trp synthetic dropout mix | Formedium | Cat#DCS0149 |
| Guanidine hydrochloride | Sigma | Cat#G4505-1KG |
| HaloTEV Protease | Promega | Cat#G6601 |
| Critical Commercial Assays |  |  |
| cOmplete EDTA-free protease inhibitor cocktail tablets | Roche | Cat#11873580001 |
| Ni-NTA Superflow | QIAGEN | Cat#30410 |
| Calmodulin Affinity resin | Agilent | Cat#214303-52 |
| Pierce spin columns snap cap | Thermo Scientific | Cat#69725 |
| Slide-A-lyzer cassette 2000MWCO | Thermo Scientific | Cat#66205 |
| RNase-It Ribonuclease cocktail | Agilent | Cat#400720 |
| RNasin Ribonuclease Inhibitor | Promega | Cat#N21150 |
| Recombinant RNasin Ribonuclease Inhibitor | Promega | Cat#N2511 |
| DNase RQ1 | Promega | Cat#M6101 |
| T4 RNA Ligase 2, truncated K227Q | NEB | Cat#M0351 |
| T4 RNA Ligase 1 | NEB | Cat#M0204L |
| T4 PNK | NEB | Cat#M0201L |
| Nitrocellulose membranes | GE Healthcare | Cat#10 439 196 |
| MetaPhor agarose | Lonza | Cat#50180 |
| NuPAGE 4-12% polyacrylamide Bis-Tris Gels | Life Technologies | Cat#NP0335 |
| NuPAGE LDS 4x sample buffer | Life Technologies | Cat#NP0007 |
| NuPAGE SDS-MOPS running buffer | Life Technologies | Cat#NP0001 |
| NuPAGE Transfer Buffer | Life Technologies | Cat#NP00061 |
| MinElute Gel Extraction kit | QIAGEN | Cat#28604 |
| Proteinase K | Roche | Cat#03115836001 |
| RNase H | NEB | Cat#M0297L |
| LA Taq | Takara | Cat#RR002M |
| Deposited Data |  |  |
| Raw data files from CRAC | NCBI Gene expression omnibus | GSE136056 |
| Raw image files | Mendeley | DOI:<br>10.17632/m253kk9sm6.1 |
| Experimental Models: Organisms/Strains |  |  |
| S. cerevisiae Strain background: BY4741 (MATa his3Δ1 leu2Δ0 met15Δ0 ura3Δ0) | <sup>69</sup> | yTWT001 |
| S. cerevisiae Strain Rpa190HTP a his3Δ1 leu2Δ0 met15Δ0 ura3Δ0 RPA190-HTP::URA3MX | This study | yTWT046 |
| S. cerevisiae Strain Rpa135 HTP a his3Δ1 leu2Δ0 met15Δ0 ura3Δ0 RPA135-HTP::URA3MX | This study | yTWT051 |
| S. cerevisiae Strain Rpa135 HTP Rpa12ΔC a his3Δ1 leu2Δ0 met15Δ0 ura3Δ0 RPA12(1-74aa only) RPA135-HTP::URA3MX | This study | yTWT232 |
| S. cerevisiae Strain Rpa190 HTP 25 rDNA a ade2-1 ura3-1 his3-11,15 trp1-1 leu2-3,112 can1-100 fob1Δ::HIS3 RPA190-HTP::URA3MX | This study | yTWT144 |
| S. cerevisiae Strain nsi1Δ Rpa135 HTP a his3Δ1 leu2Δ0 met15Δ0 ura3Δ0 RPA135-HTP::URA3MX nsi1ΔHIS | This study | D1436 |
| S. cerevisiae Strain nsi1Δ Rpa135 HTP Rpa12ΔC a his3Δ1 leu2Δ0 met15Δ0 ura3Δ0 RPA12(1-74aa only) RPA135-HTP::URA3MX nsi1ΔHIS | This study | D1433 |
| S.cerevisiae strain Rat1-TAP::HIS a his3Δ1 leu2Δ0 met15Δ0 ura3Δ0 Rat1-TAP::HIS | Open biosystems |  |
| S.cerevisiae strain Rai1-TAP::URA a his3Δ1 leu2Δ0 met15Δ0 ura3Δ0 Rai1-TAP::URA | This study | D1375 |
| S cerevisiae strain Trf4-TAP a his3D1 leu2D0 met15D0 ura3D0 TRF4-TAP-his | Open biosystems |  |

| Software and Algorithms |  |  |
| --- | --- | --- |
| PyCRAC | Webb et al., 2014 | https://bitbucket.org/sgrann/pycrac |
| SAMtools v1.3.1 | Li et al., 2009 | http://www.htslib.org/ |
| Bedtools v2.25 | <sup>70</sup> | https://github.com/arq5x/bedtools2 |
| Prism 7 | Graphpad | http://www.graphpad.com |
| Integrative Genomics Viewer | Broad Institute | http://software.broadinstitute.org/software/igv/ |
| STAR v2.7.10a | PMID: 23104886 | n/a |
| trxtools 0.2.1 | This study | https://github.com/TurowskiLab/trxtools |

### EXPERIMENTAL MODEL and SUBJECT DETAILS

#### Strains

Yeast analyses were performed in strains derived from BY4741 (*MAT*a; *his3Δ1*; *leu2Δ0*; *met15Δ0*; *ura3Δ0*). For CRAC analyses, cells were grown in synthetic medium with 2% glucose at 30°C. Oligonucleotides used for strain construction are listed in Supplementary Table S1.

### METHODS DETAILS

#### *In-vivo* RNA crosslinking

Strains for CRAC experiments were grown in SD medium with 2% glucose, lacking tryptophan to OD_600_=0.5. Actively growing cells were cross-linked in culture media using megatron UVC cross-linker ^71^ for 100 seconds or the VariX cross-linker for 8 seconds. Cells were span and washed with 1x PBS buffer before freezing at -80°C.

#### CRAC

Samples were processed as previously described ^49^. However, phosphatase treatment was omitted, so the 3’-OH ends required for linker ligation are present only on nascent RNA transcripts. Cells were lysed in TNC100 (50mM Tris-HCl pH7.5, 150mM NaCl, 0.1% NP-40, 10mM CaCl_2_, 5mM β-mercaptoethanol, 50U of DNase RQ1 and a protease-inhibitor cocktail (1 tablet / 50mL) with zirconia beads in a 50mL conical. The cells were lysed with five one-minute pulses, with cooling on ice in between. The supernatant was spun for 20 minutes at 21,000g. The cleared lysate was incubated with the IgG Sepharose for two hours at 4°C, with nutating. Subsequently, the beads were washed three times with TN1000 (50mM Tris-HCl pH7.5, 1000mM NaCl, 0.1% NP-40) and two times TN100 (50mM Tris-HCl pH7.5, 100mM NaCl, 0.1% NP-40). Protein:RNA complexes were eluted by incubation with HaloTEV for 2h at 18°C with shaking. The eluate was transferred to a fresh tube, 2.5U of RNace-IT was added and samples were incubated for 5 minutes at 37°C to fragment protein-bound RNA.

The 500 μL eluate was adjusted for nickel affinity purification with the addition of 400 mg guanidine hydrochloride, 45μL NaCl (3M) and 3μL imidazole (2.5M) and added to 100μL of washed nickel beads.

Following overnight incubation, the nickel beads were washed three times with WBI (6.0M guanidine hydrochloride, 50mM Tris-HCl pH7.5, 300mM NaCl, 0.1% NP-40, 10mM imidazole and 5mM β-mercaptoethanol), three times with PNK buffer (50mM Tris-HCl pH7.5, 50 mM NaCl, 1.5mM MgCl2, 0.1% NP-40, and 5mM β-mercaptoethanol) and transferred to a spin column.

Subsequent reactions (80μL total volume for each) were performed in the columns, and afterward washed once with WBI and three times with PNK buffer:

1. 3’ linker ligation (1x PNK buffer (NEB), 10% PEG8000, 20U T4 RNA Ligase II truncated K227Q, 80U RNasIN, 80pmol pre-adenylated 3’ miRCat-33 linker (IDT); 24°C 6hrs).
2. 5’ end phosphorylation and radiolabeling (1x PNK buffer (NEB), 40U T4 PNK (NEB), 40µCi ^32^P-γATP; 37°C for 60min, with addition of 100nmol of ATP after 45min).
3. 5’ linker ligation (1x PNK buffer (NEB), 10% PEG8000, 40U T4 RNA ligase I (NEB), 80U RNasIN, linker, 200pmol 5’ linker, 1mM ATP; 16°C, overnight).

The beads were washed once with WBI and three times WBII (50mM Tris-HCl pH7.5, 50mM NaCl, 0.1% NP-40, 200mM imidazole, and 5mM β-mercaptoethanol) buffer. Protein:RNA complexes were eluted in 400μL of elution buffer (50mM Tris-HCl pH7.5, 50mM NaCl, 0.1% NP-40, 200mM imidazole, and 5mM β-mercaptoethanol) and TCA precipitated for 1hrs on ice. RNPs were pelleted at 21,000g for 20min, washed in cold acetone and resuspended in 30μL 1X NuPAGE sample loading buffer supplemented with 8%β-mercaptoethanol. The sample was denatured by incubation at 65°C for 5min, and run on a 4%–12% Bis-tris NuPAGE gel at 130V. The protein:RNA complexes were transferred to Hybond-C nitrocellulose membranes with NuPAGE MOPS transfer buffer for 2h at 100V.

Labelled RNA was detected by autoradiography. The appropriate region was excised from the membrane and treated with 0.2μg/μL Proteinase K (50mM Tris-HCl pH7.5, 50mM NaCl, 0.5%SDS, 1mM EDTA; 2hr 55°C with shaking) in a 400μL reaction. The RNA component was isolated with a standard phenol:chloroform extraction followed by ethanol precipitation with 1µL of GlycoBlue. The RNA was reverse transcribed using Superscript III and the miRCat-33 RT oligo (IDT) for 1h at 50°C in a 20μL reaction. The resulting cDNA was amplified by PCR in 50µL reactions using La Taq (5 μL template, 21-26 cycles) PCR reactions were combined, precipitated in ethanol, and resolved on a 3%Metaphore agarose gel. A region corresponding to 140 to 200 bp was excised from the gel and extracted using the Min-elute kit. Libraries were measured with Qbit and sequenced using Illumina HiSeq or llumina MiniSeq with 75bp single-end reads.

#### Tap purification of Rat1, Rai1 and Trf4

We attempted to co-express tagged Rat1 and Rai1 in *E.coli* and purified the complex ^39,72–74^, but due to low activity of the exonuclease we used Rat1-Rai1 complex purified from yeast. Strains were grown in YPD to OD 1.5, harvested, washed with ice cold PBS and frozen at -80°C. Lysis was performed in TMN100 buffer as previously described. Lysates were bound to IgG-sepharose for 1.5 h, washed 3 times in TMN100 and eluted with Halo-TEV at 24°C for 2 h. TEV eluate was bound to Calmodulin resin for 2 h at 4°C, washed in 3 times TMN150/2 mM CaCl2 and eluted with 5mM EGTA, 10m M Tris pH 8, 50 mM NaCl. 500μL of eluate were injected into a Slide-A-lyzer cassette and dialyzed for 3h TMN75 / 50% glycerol.

#### Purification of RNA polymerase I and in vitro assay

Strains were grown in YPD to OD_600_=1.5-2. Cells were span and washed with 1x PBS buffer before freezing at -80°C.

Cells were lysed in Lysis buffer (50mM Hepes pH7.8, 400mM (NH_4_)_2_SO_4_,10% glycerol, 40mM MgCl_2_, 3mM DTT, 10mM CaCl_2_, 50U of DNase RQ1 and a protease-inhibitor cocktail (1 tablet / 50mL) with zirconia beads in a 50mL conical. The cells were lysed with ten one-minute pulses, with cooling on ice in between. The supernatant was spun for 20min at 21,000g.

The protein content of the supernatant was determined using the Bradford assay. Equal protein amounts (usually 1ml cell extract, 20–30 mg) were incubated with 50–75μl of immunoglobulin-G (rabbit IgG, I5006, Sigma) coupled magnetic beads slurry (Dynabeads M-270 Epoxy, 300mg) for 1–2 h on a rotating wheel. The beads had previously been equilibrated with lysis buffer. The beads were washed four times with 1 ml buffer B1500 (20mM HEPES/KOH pH7.8, 1.5 M KOAc, 1mM MgCl_2_, 10% glycerol, 0.1% IGEPAL CA-630) and three times with 1ml buffer B200 (20mM HEPES/KOH pH7.8, 200mM KOAc, 1mM MgCl_2_, 10% glycerol). For elution, beads were finally resuspended in 400μl of TMN150 buffer (50mM Tris pH7.8, 150mM NaCl, 1.5mM MgCl_2_,0.1% NP40, 5mM BetaMEth), supplemented with 3μl TEV protease (HaloTEV, Promega G6602) and incubated for 2h at 24°C in a thermomixer (1,000 rpm). The supernatant was collected, glycerol was added to 10% and aliquots were stored at -20°C for short term or at −80°C for longer. For buffer exchange assays, TEV elution was omitted, and aliquots were stored only for short term at 4°C. 10% of the purified fraction was analyzed via SDS–PAGE to monitor the purification success. Protein concentrations were determined by comparing the intensity of Coomassie-stained RNA polymerase subunits to the defined amount of Coomassie-stained HaloTEV protease used.

The *in vitro* RNA extension assay was modified from ^26,75^. For 1 reaction, 2pmol of annealed RNA-DNA-DNA scaffold was preincubated with ∼2pmol of purified enzyme for 20min at 20°C. Transcription was started by adding 6μl-2x transcription buffer (TB). Elongation was performed in 1x TB (60mM (NH4)2SO4, 20mM HEPES/KOH pH 7.6, 8mM MgSO4, 10µM ZnCl_2_, 10% glycerol, 10mM DTT) supplemented with 1mM NTPs. The samples were incubated at 28°C for 5min. For backtracking assays, reaction tubes were placed on a magnetic rack, and supernatant was removed. Beads were washed with 200μl buffer B200, resuspended in 12μl 1x TB without NTPs and incubated at 28°C for 10min. All reactions were stopped by addition of 2x RNA loading dye (Thermo, R0641). Samples were heat denatured at 80°C for 5min and resolved on 8M urea 20% polyacrylamide gels. Fluorescently labeled transcripts were visualized using a FujiFilm FLA-5100 Fluorescent Image Analyser. Images were processed using Multi Gauge software (Fuji). Oligonucleotides used for in vitro assays are listed in Table S2.

### Polyadenylation assays

For 1 reaction, 2pmol of annealed RNA-biotinDNA-DNA scaffold bound to Streptavidin magnetic beads was preincubated with ∼2pmol of purified enzyme for 20 min at 20°C. Beads were washed twice with B200 buffer. Transcription was started by adding 6μl 2x transcription buffer (TB). Elongation was performed in 1xTB (60mM (NH_4_)_2_SO_4_, 20mM HEPES/KOH pH 7.6, 8 mM MgSO_4_, 10µM ZnCl_2_, 10mM DTT) supplemented with 1mM NTPs-GTP containing 0.5μl [alpha-^32^P]-ATP at 28°C for 10 min. Beads were washed 3 times with B200 and resuspended in 1xTB. ∼2pmol of purified Trf4,1mM ATP were added and timepoints were taken at point 0, 5, 15 and 30minutes. Poly(A)-tails were analyzed on 15% Acrylamide/Urea gels.

### Termination assays

For the scaffold of the termination assays 5’ kinased input RNA was used.

For 1 reaction, 2pmol of annealed RNA-DNA-DNAbiotin scaffold bound to Streptavidin magnetic beads was preincubated with ∼2pmol of purified enzyme for 20min at 20°C. Beads were washed twice with B200 buffer. Transcription was started by adding 6μl 2x transcription buffer (TB). Elongation was performed in 1x TB (60mM (NH_4_)_2_SO_4_, 20mM HEPES/KOH pH 7.6, 8mM MgSO_4_, 10µM ZnCl_2_, 10mM DTT) supplemented with 1mM NTPs-GTP containing 0.5μl 32P-alpha-ATP at 28C for 10 minutes. Beads were washed 3 times with B200 and resuspended in 1xTB. ∼2pmol of purified Rat1/Rai1 were added and timepoints were taken at point 0, 5, 15,30 and 60 minutes. PolyA-tails were analyzed on 15% Acrylamide/Urea gels. To follow the protein drop-off rate, the termination assay was performed omitting the 32P-alpha-ATP. Samples were analyzed on per-cast SDS 4-12% gradient gels and westerns were done with TAP-antibody. Drop off rate was calculated using ImageJ.

### Modification of Mathematical model for RNAPI transcription

The numerical model to estimate premature termination was developed on the basis of previously developed model ^7^ as described in the main text.

#### Simulation conditions

Most of the simulation conditions was used as previously: gene length 7000 [nt], RNAPI size 38 [nt], RNA-DNA hybrid within the transcription bubble 11 [nt], transcription initiation probability 0.8, average RNAPI velocity 50 [nt·s^-1^]. Model parameters: 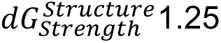, 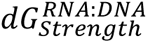 which was represented as a ratio to 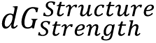, with ratio = 0.48, structure2consider = -11. Time step was 0.004 [s].

To simulate a full division cycle, we extended simulation time for 6000 [s] (=100 min) and run parallel simulations (n=16). To calculate output of 75 transcription units (∼50% active out of 150) we multiplied the output by 4.7.

Given alternative models we tested following parameter combinations: DNA stiffness constant c = {0,500},: 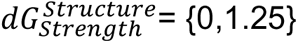 premature termination probability = {0, 0.01, 0.02, 0.03, 0.04, 0.05, 0.06, 0.07, 0.08, 0.09, 0.1, 0.2, 0.3, 0.4, 0.5, 0.6, 0.7, 0.8, 0.9, 1}i, premature termination distance = {2000, 6750}. This gives us a total of 226 sets of parameters which were subsequently evaluated to ensure that the chosen parameter set gives a representative data, approximately in the middle of the set of simulations with neighboring parameters. Full set of parameters was uploaded to the repository as CSV file runParameters.csv https://github.com/tturowski/RNAPI-model-v2. Output data were saved in MAT format and analyzed using jupyter-notebook uploaded to https://github.com/tturowski/Pol1_termination_MS as Fig_6_model_output.ipynb.

### QUANTIFICATION AND STATISTICAL ANALYSIS

#### Analysis reproducibility

To support transparency and reproducibility of the analysis all steps were performed using snakemake pipeline and jupyter-notebooks. The files were deposited in git repository https://github.com/tturowski/Pol1_termination_MS.

#### Pre-processing and data alignment

Illumina sequencing data were demultiplexed using in-line barcodes and in this form were submitted to GEO. First quality control step was performed using FastQC software (http://www.bioinformatics.babraham.ac.uk/projects/fastqc/) considering specificity of CRAC data.

All processing steps were performed using snakemake pipeline (v7.14.0 [34035898]). Raw reads were collapsed to remove PCR duplicates using FASTX-collapser v0.0.14 (http://hannonlab.cshl.edu/fastx_toolkit/) then inline barcodes were removed using FASTX-trimmer v0.0.14. The 3’ adapter were removed using flexbar v3.5.0 ^76^ with parameters - ao 4 –u 3 -m 7 -n 4 -bt RIGHT, and filtered to retain only reads containing the 3’ adaptor. All datasets were aligned to the yeast genome (sacCer3, EF4.74) using STAR v2.7.10a and saved as bam file format. Bam files were sorted and indexed using samtools v1.15.1 ^77^. Second quality control step was performed using featureCounts v2.0.1 which calculates overlaps between aligned cDNAs and yeast genomic features. BigWig files were generated using bamCoverage script from deepTools package v3.5.1 ^78^.

#### Selection of the 5’ ends, 3’ ends and polyA

To prepare BigWig files containing the 5’ ends of reads, or the 3’ ends of reads, or the 3’ ends of poly-adenylated reads (AAA or longer) custom tool SAM2profilesGenomic.py from trxtools v0.2.2 was used. The package is publicly available https://pypi.org/project/trxtools/0.2.2/.

#### RNA polymerase I profile

Downstream analyses were performed using python 2.7 Jupiter notebooks, python libraries (pandas v0.19.2, numpy v1.16.0, scipy v1.2.0, matplotlib v2.2.3) and in-house functions available in trxtools, created as modification of gwide toolkit published previously^49^. All reads mapping to the gene encoding pre-rRNA (*RDN37* gene with 1300 nt overhangs) were summed up to 1 and fraction of reads was used further, adding 10^-7^ pseudo count. There are two copies of the *RDN37* gene in the reference genome; *RDN37-1* and *RDN37-2.* Subsequent analyses used the combination of *RDN37-1* and *RDN37-*2 gene or *RDN37-2* for analysis of transcriptional read-through.

The smooth data we used centered Blackman function (window 10). CRAC profiles were presented similar to boxplots of six biological replicates: median as a solid line, range between second and third quartile with darker color and range between minimum and maximum as lighter color. The basic profile of RNAPI CRAC was established as previously described ^7^.

### Peak calling and metaplots

Peak calling was performed using findPeaks function from trxtools v0.2.2 using order value 45 and window 80. To generate peak metaplot for each peak or trough, a two sided window around the feature was superimposed with all other peaks. Mean for all windows were calculated and data for each dataset were presented as peak metaplot. Metaplots were generated using cumulativePeaks function from trxtools v0.2.2.

### Statistical analyses and numerical methods

Most of the plots and statistical analyses of this work were performed using python 3.6, Jupiter notebooks and python library scipy v1.2.0. Boxplots present 2^nd^ and 3^rd^ quartile, line marks median and whiskers range between 5^th^ and 95^th^ percentile. To visualize reproducible differences between strains we marked area between outside q2-q3 range.

The Western blot quantification was conducted using ImageJ software. To determine the RNAPI drop-off rate, the percentage change between the 0-minute time point and the given time point was calculated as a ratio and then 100% was subtracted. This adjustment normalizes the 0-minute time point to 0%, and the result represents the percentage of drop-off for subsequent time points.

Statistical significance was assessed using Wilcoxon signed-rank and rank sum tests, with the T-test applied where appropriate (indicated in figure legend). All statistical tests were two-tailed, and the legend for p-values was consistently represented as follows: * p<0.05, ** p<0.01, *** p<0.001.

### RESCOURCE AVAILABILITY

#### Lead Contact

Further information and requests for resources and reagents should be directed to and will be fulfilled by the Lead Contact, David Tollervey.

#### Materials Availability

All unique/stable reagents generated in this study are available from the Lead Contact without restriction.

#### Data and Code Availability

All RNA sequencing data from this study have been submitted to the NCBI Gene Expression Omnibus (GEO; http://www.ncbi.nlm.nih.gov/geo/) and are available under accession number GSE246546.

The full MATLAB code for the mathematical model has been submitted as a git repository: https://github.com/tturowski/RNAPI-model-v2

The analysis steps were deposited in git repository https://github.com/tturowski/Pol1_termination_MS

## SUPPLEMENTARY MATERIAL

**Figure S1.**
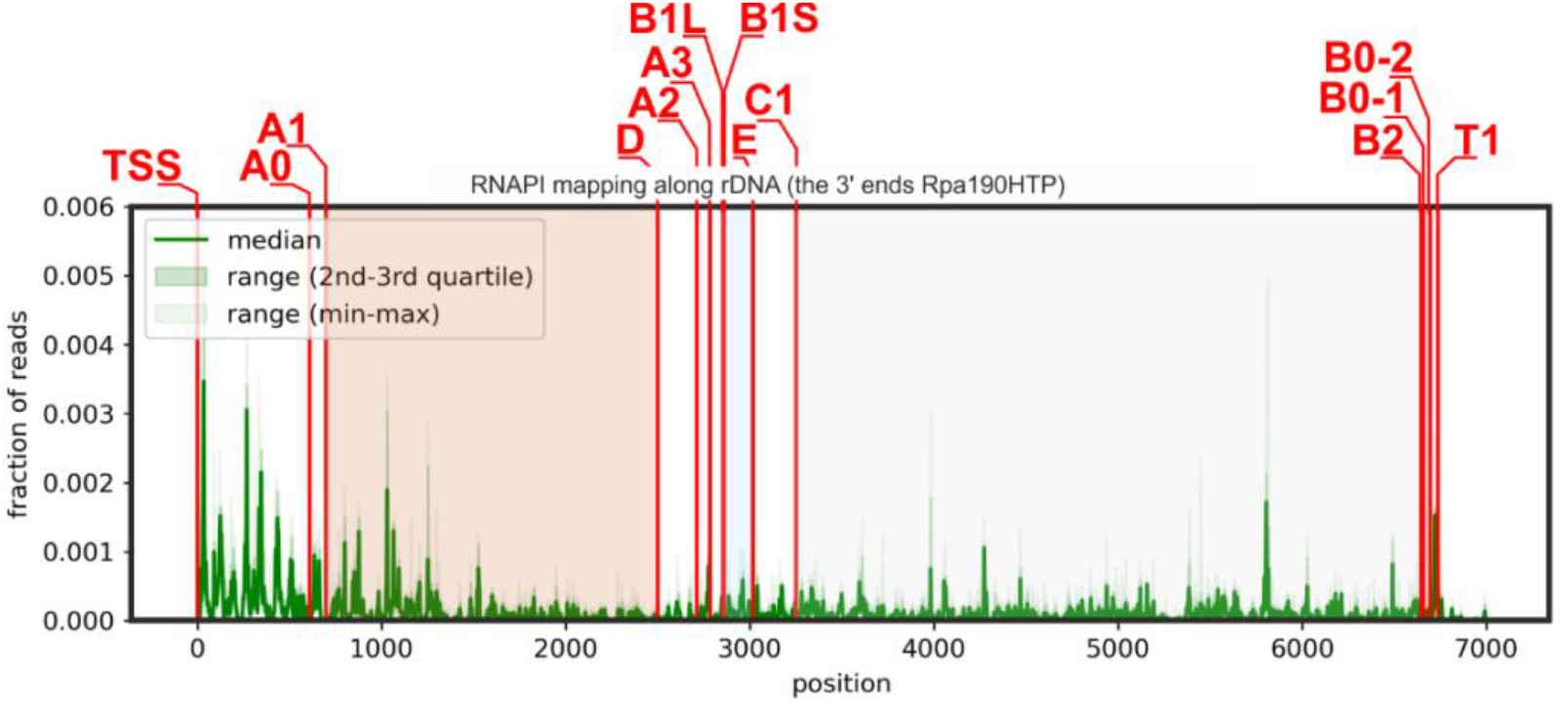
Scheme of yeast rDNA transcription unit with marked processing sites.

**Figure S2.**
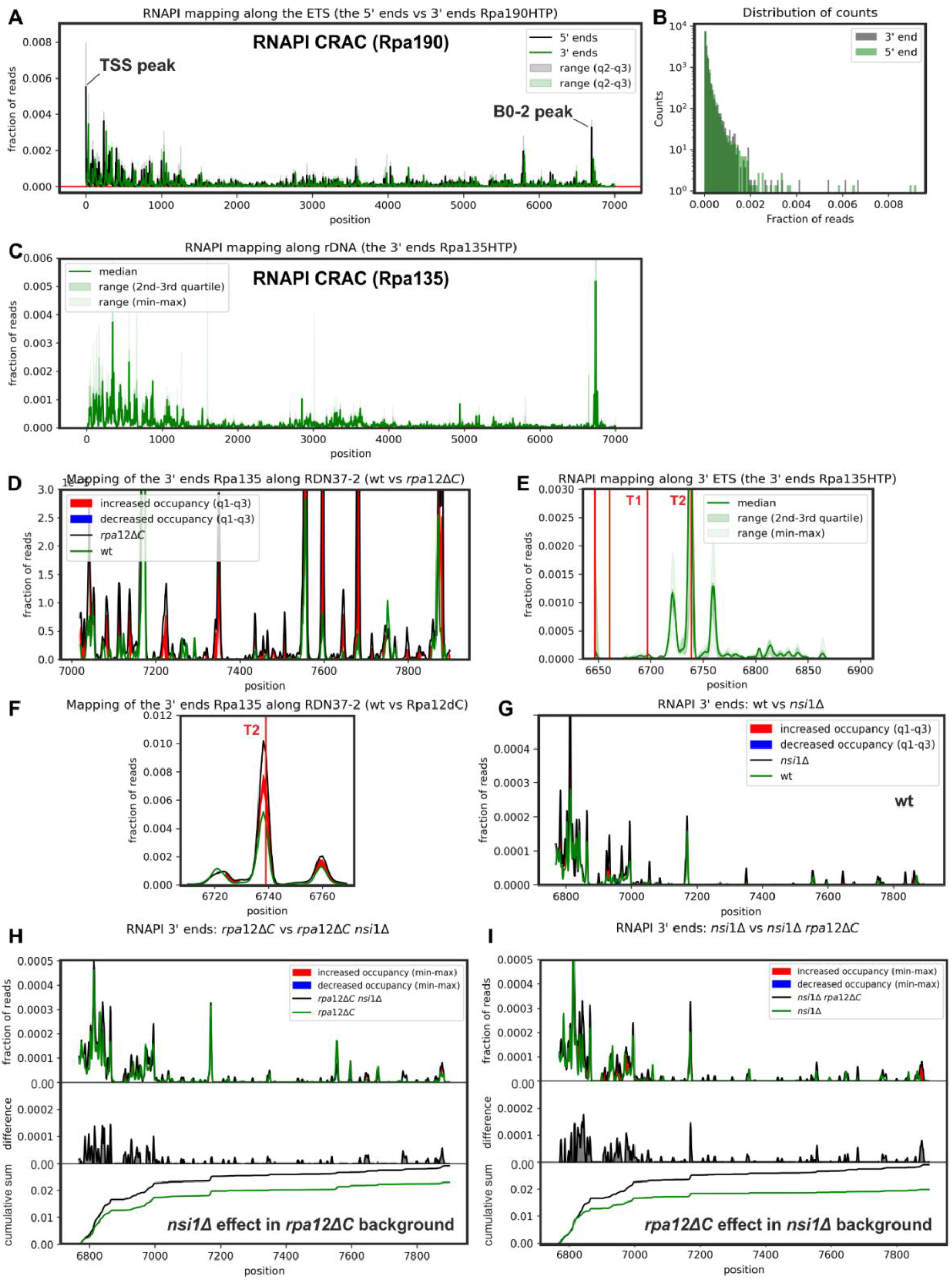
RNAPI termination is associated with decreased elongation kinetics. A: RNAPI Rpa190-HTP CRAC mapping of the 5’ ends (black) and the 3’ ends (green) of reads. Note that substantial differences between the 5’ and 3’ ends are visible for the very 5’ end at the transcription start site (TSS peak) and within the 3’ ETS, at the B0-2 cleavage site (B0-2 peak). B: 5’ ends of RNAPI CRAC reads are less distributed relative to 3’ ends. C: Distribution of RNAPI Rpa135-HTP CRAC 3’ ends of reads along the rDNA. D: Transcriptional read-trough is increased in *rpa12ΔC* strain in comparison to wt. E: Distribution of RNAPI Rpa135-HTP CRAC reads around the 3’ end of the 35S pre-rRNA. F: RNAPI Rpa135-HTP CRAC signal is increased at the T2 site in Rpa12ΔC strain. G: Limited effect of *nsi1Δ* on RNAPI transcriptional read-trough detected with Rpa135-HTP CRAC in wt background. H: Limited effect of *nsi1Δ* on RNAPI transcriptional read-trough detected with Rpa135-HTP CRAC in *rpa12ΔC* background. I: Effect of *rpa12ΔC* on RNAPI transcriptional read-trough detected with Rpa135-HTP CRAC in *nsi1Δ* background.

**Figure S3.**
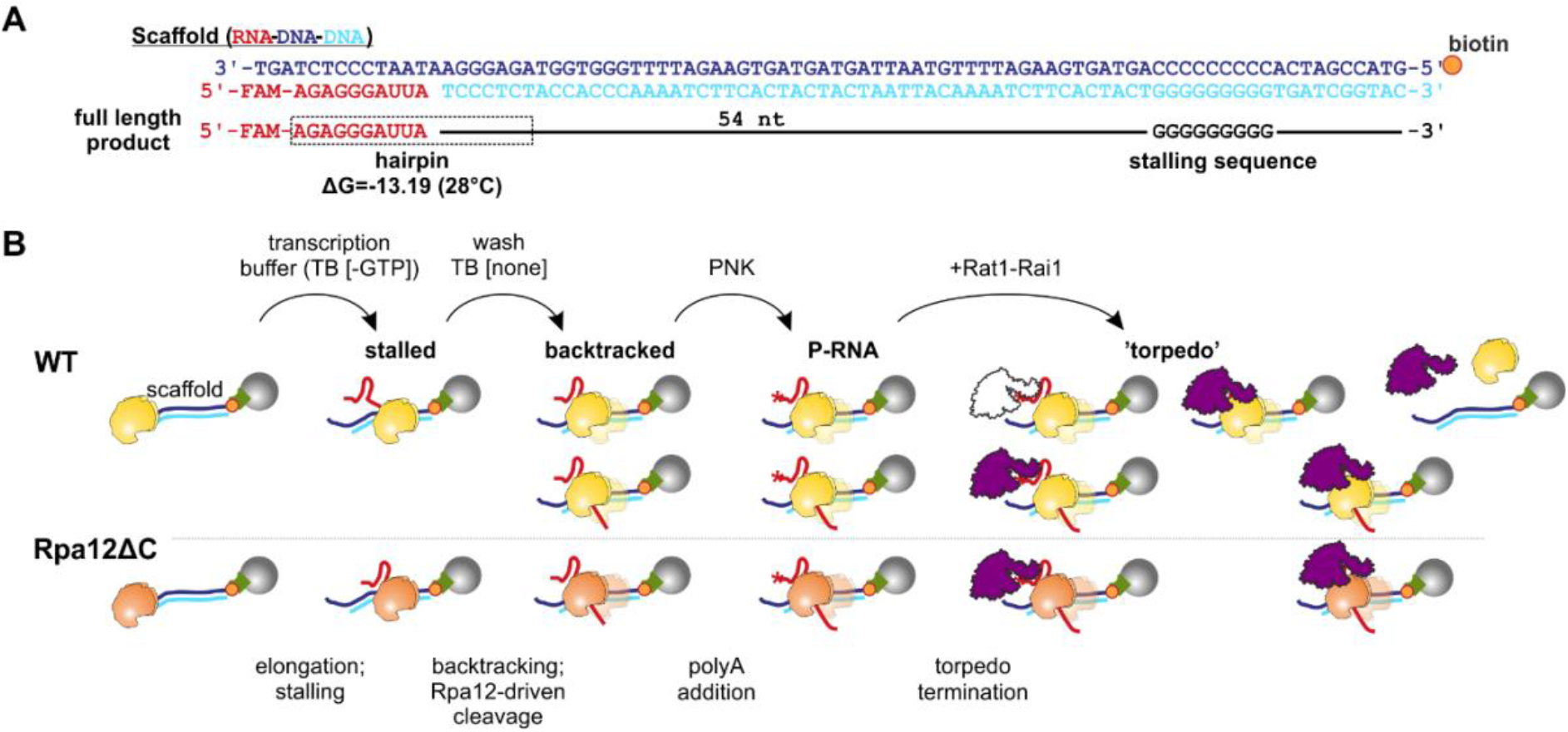
Schematic of *in vitro* termination assay. A: Sequence of RNA-DNA-DNA scaffold immobilized via biotin-streptavidin interaction. B: Schematic of the experiment.

**Figure S4.**
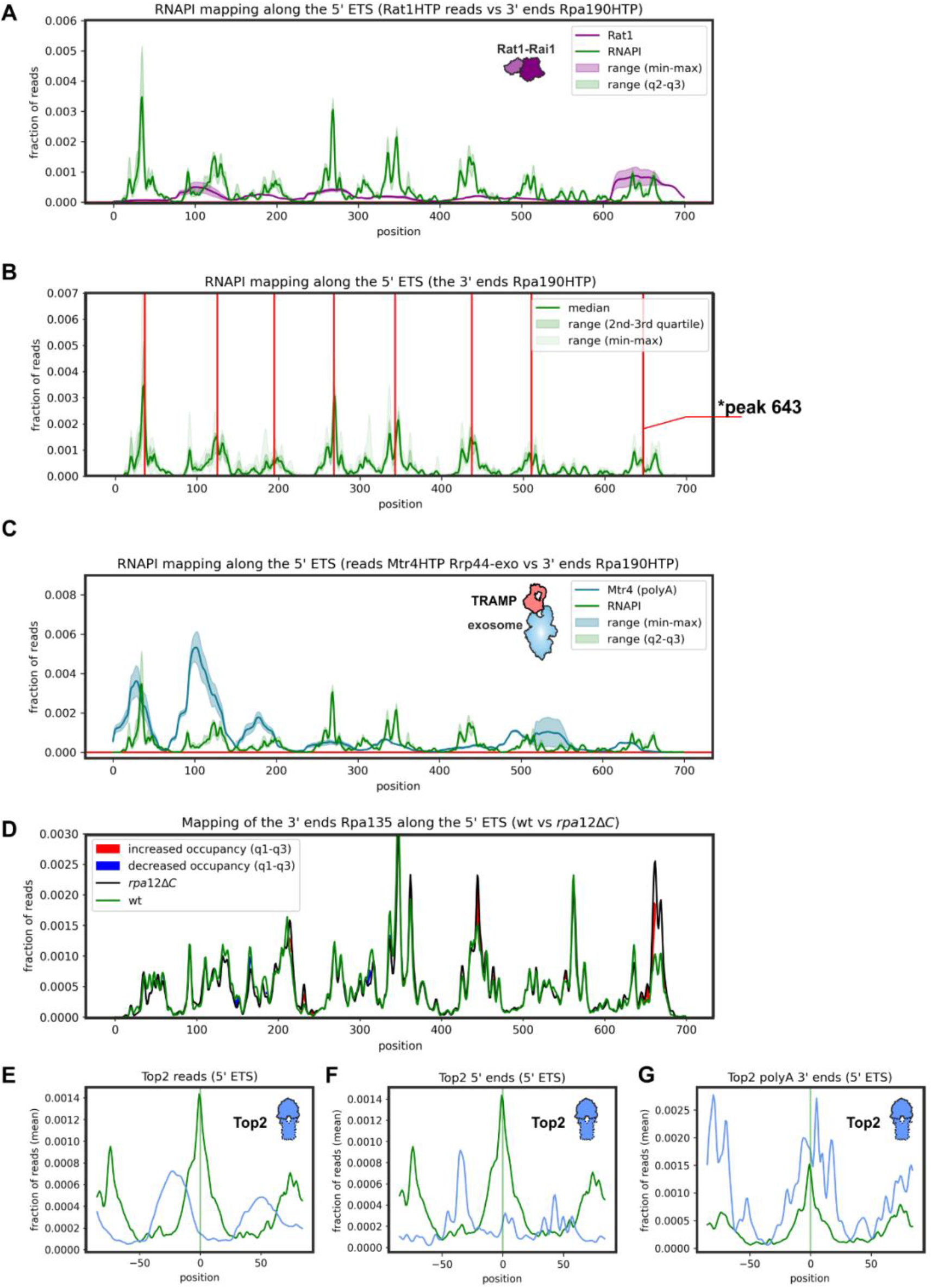
Rat1 and TRAMP are found at sites of slowed RNAPI elongation. A: Rat1 CRAC reads superimposed with the 3’ ends of RNAPI CRAC reads. B: Peaks found with peak-calling algorithm that were used to generate peak metaplot. * Note peak 643 was excluded from meta-analysis to separate Rat1 surveillance from canonical degradation of the 5’ ETS during pre-rRNA processing. C: Mtr4 (Rrp44-exo) CRAC reads superimposed with the 3’ ends of RNAPI CRAC reads. D: RNAPI (Rpa135-HTP) CRAC shows minor difference in *rpa12ΔC* strain in the 5’ ETS. E-G: RNAPI Rpa190-HTP CRAC peak metaplot for the 5’ ETS, comparing Rpa190 and Top1-HTP CRAC at the level of single peaks: reads (E), the 5’ ends of reads (F) and the 3’ ends of polyadenylated reads (G).

**Figure S5.**
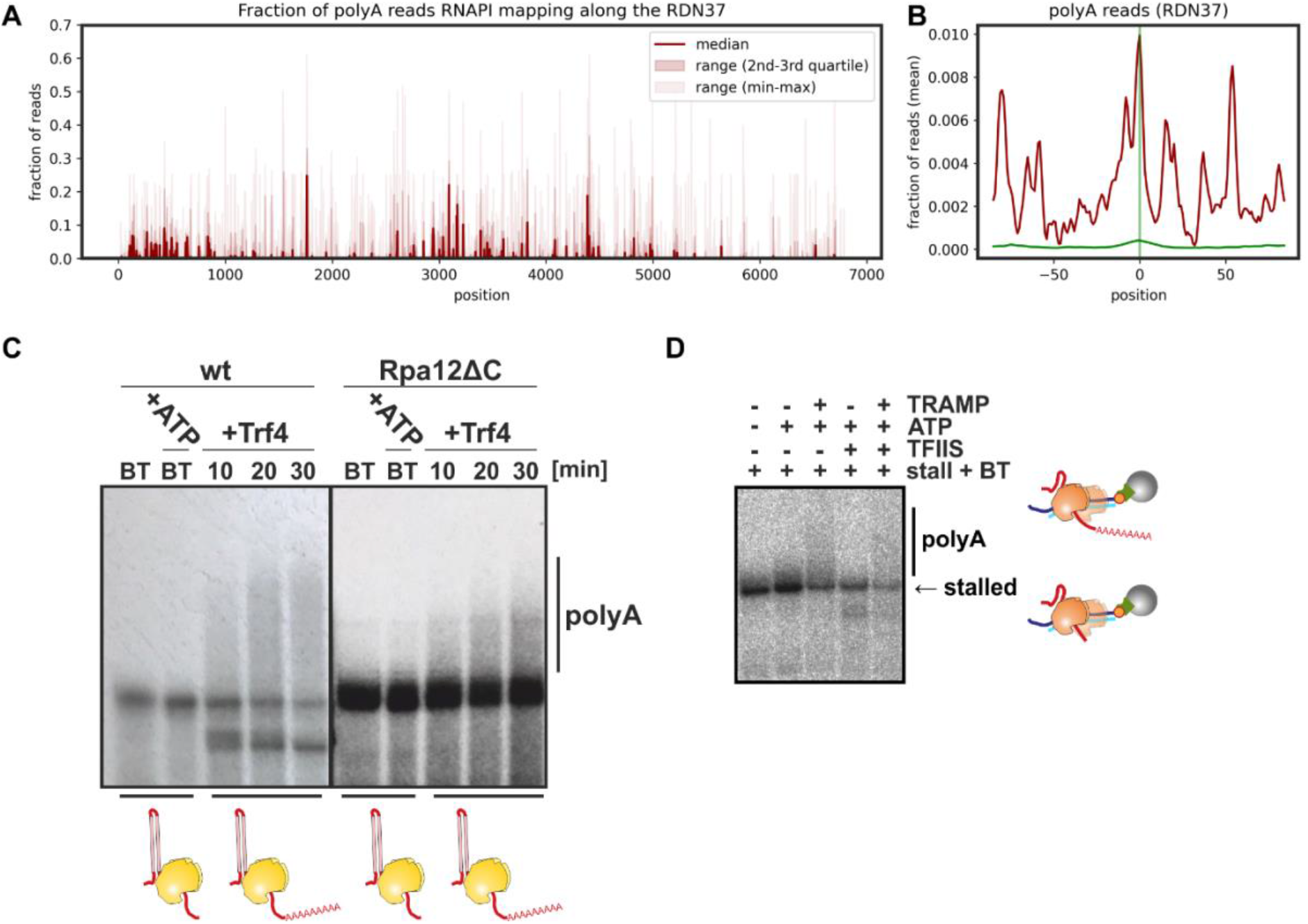
Backtracked nascent transcripts can be oligo-adenylated by the TRAMP complex A: Fraction of polyA reads mapping along the *RDN37*. B: RNAPI CRAC peak metaplot for *RDN37*, comparing the 3’ ends of the reads (green) with poly(A) reads. C-D: Trf4 oligo-adenylates the 3’ end of backtracked, nascent RNA *in vitro* extruded from RNAPI (C) and RNAPII (D).

**Figure S6.**
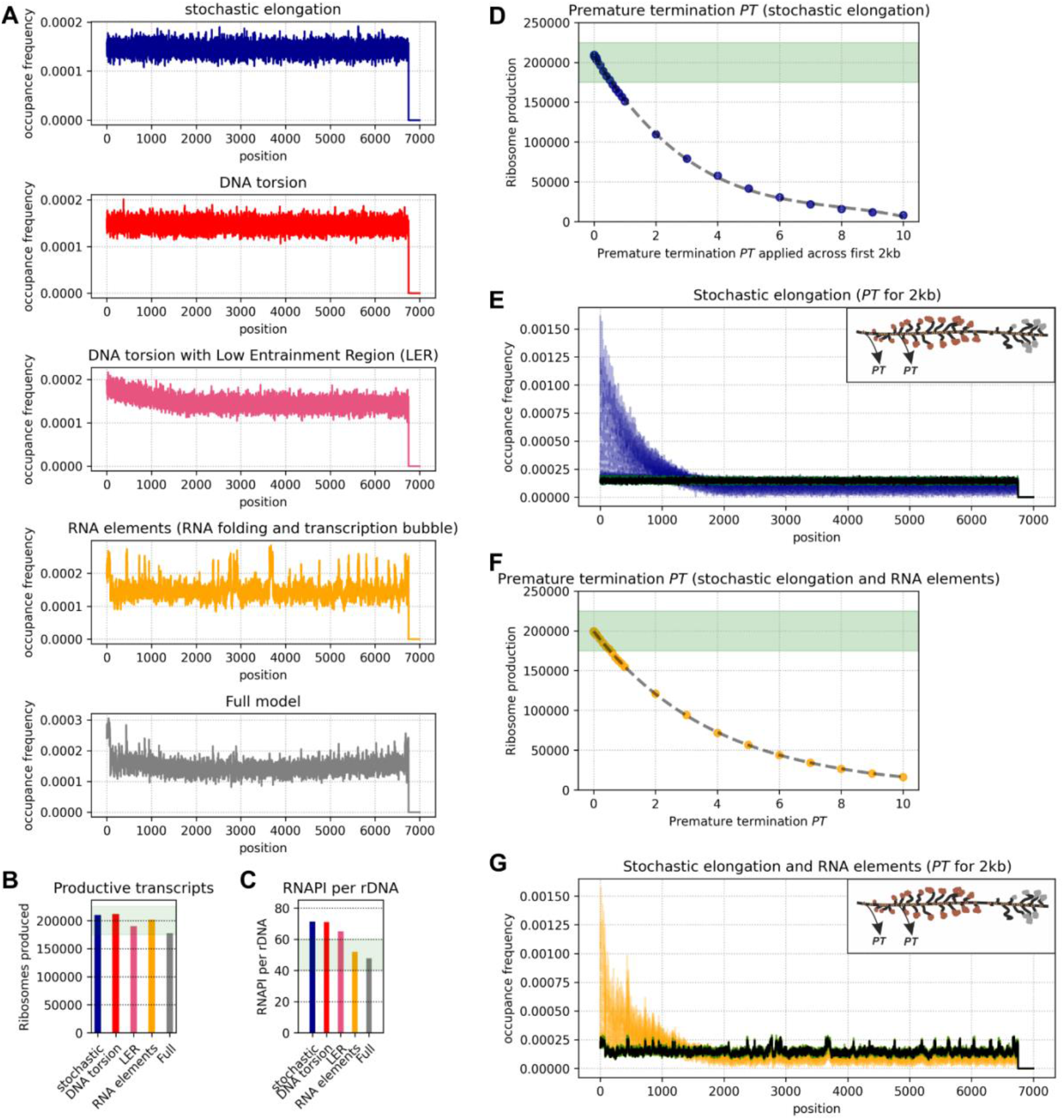
Development of premature termination *PT* function. A: Modeled RNAPI occupancy along the transcription unit using a model of stochastic initiation and discrete, stochastic elongation. The average RNAPI occupancy for 16 simulations is presented. Each simulation was run for 6,000 sec and 200 time points were collected. B: Average number of productive transcription events for each type of simulation (corresponding to panel A). C: The average number of RNAP molecules per transcription unit (corresponding to panel A). D: *PT* effect on ribosome production in model including stochastic elongation only. *PT* was applied across the first 2kb of the transcription unit. Note that because *PT* is applied only for this part of the transcription unit the values of *PT* are significantly higher than for panel Fig 6A. E: RNAPI occupancy profiles corresponding to panel D. Black – no premature termination *PT*, green *PT* where ribosome synthesis is effective, blue – ribosome synthesis is decreased or insufficient. F: *PT* effect on ribosome production in model including stochastic elongation and RNA elements. Note that because *PT* is applied only for the part of the transcription unit the values of *PT* are significantly higher than for panel Fig. 6A. G: RNAPI occupancy profiles corresponding to panel E. Black – no premature termination *PT*, green *PT* where ribosome synthesis is effective, orange – ribosome synthesis is decreased or insufficient.

**Table S1.**
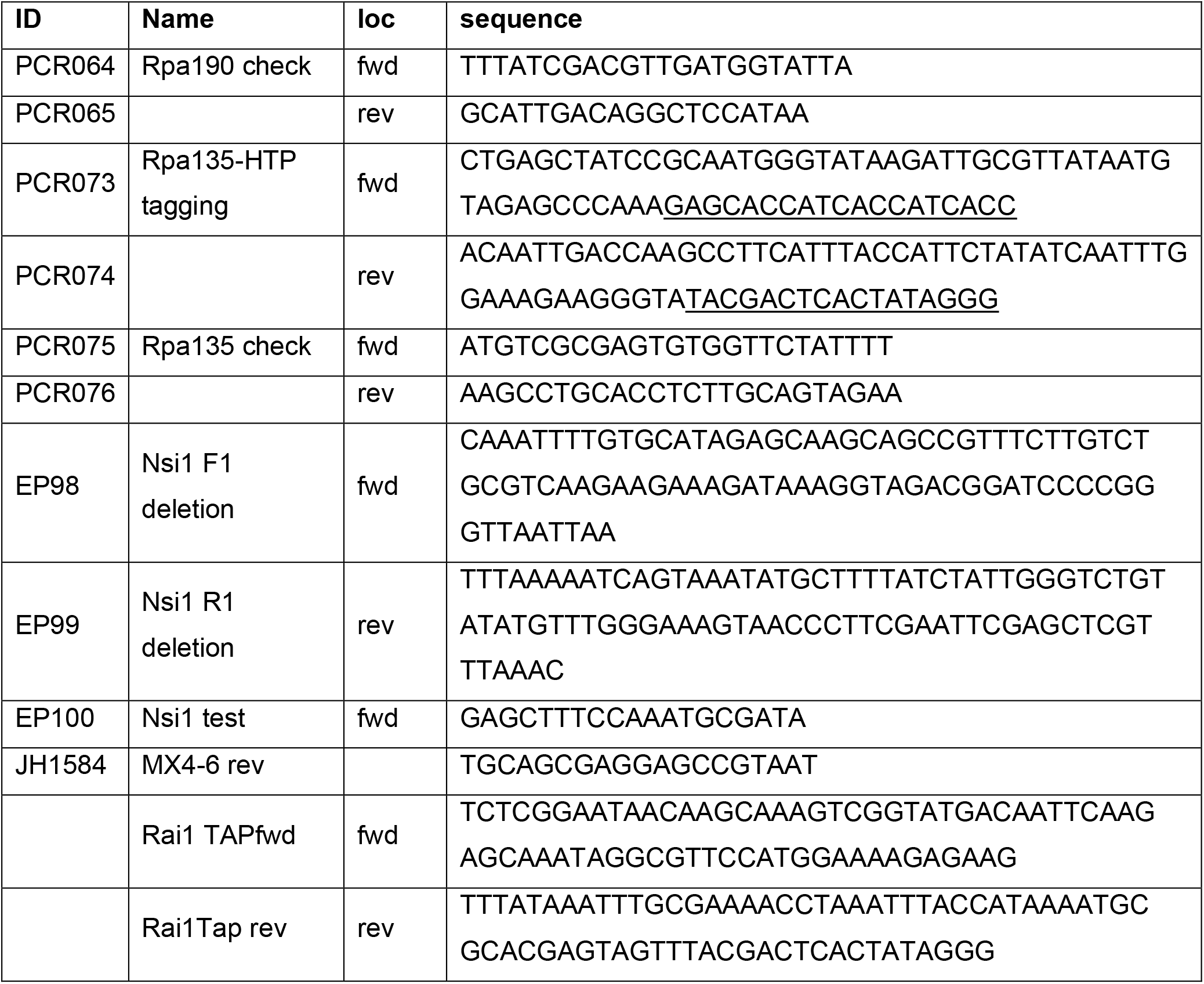
Oligonucleotides used for strain construction.

**Table S2.**
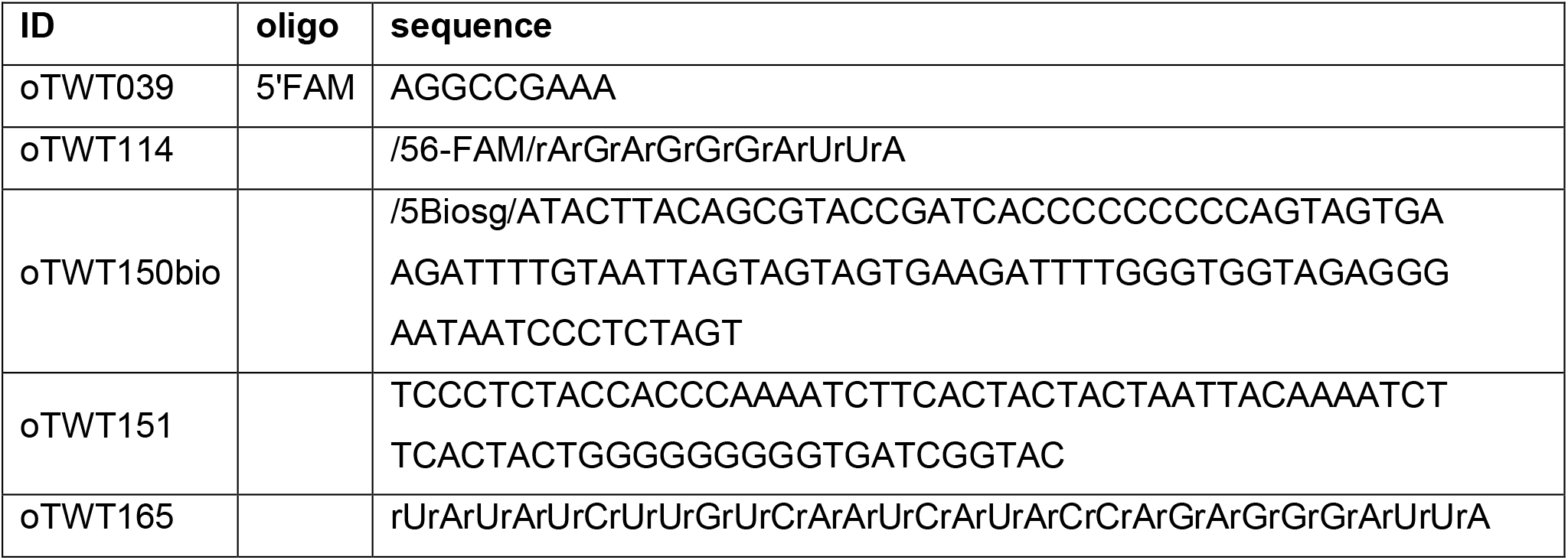
Oligonucleotides used for *in vitro* assay.

## Notes

https://github.com/tturowski/Pol1_termination_MS

### Competing Interest Statement

The authors have declared no competing interest.

