## Supplementary Informations for "Transcription termination by RNA polymerase I"

**Turowski et al**

**SUPPLEMENTARY MATERIAL**

**Figure S1.** Initial analysis of RNAPI CRAC data (related to Figure 1)
**Figure S2.** RNAPI density correlates with features in the nascent pre-rRNA (related to Figure 2)
**Figure S3.** Nascent RNA limits backtracking proportionally to folding energy (related to Figure 3)
**Figure S4.** Mathematical model of RNAPI transcription (related to Figure 4)
**Figure S5.** Conclusions from the model on the role of 5' ETS structure (related to Figure 5)
**Figure S6.** Folding of the nascent transcript plays a role in determining elongation rate of all eukaryotic RNA polymerases (related to Figure 6)

**Table S1.** Oligonucleotides used for strain construction.

**Table S2.** Oligonucleotides used for *in vitro* assay.

**Figure S1**

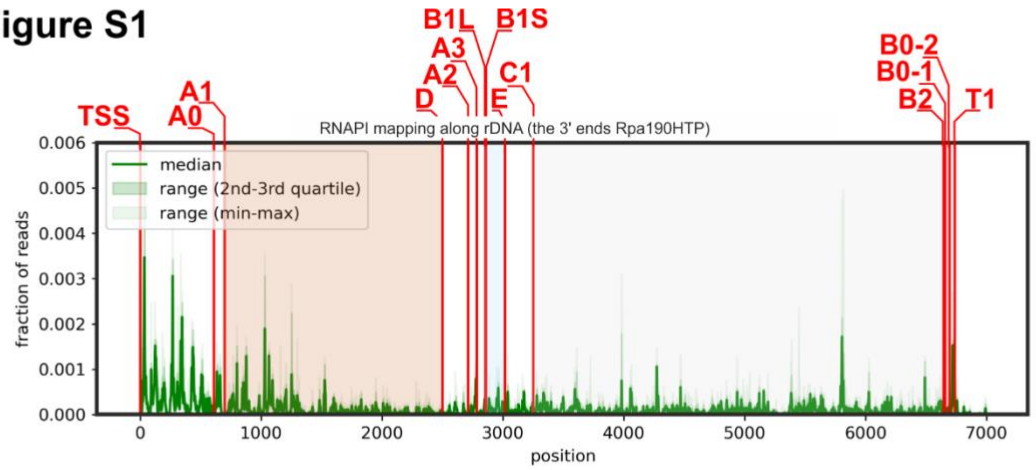


**Figure S1.** Scheme of yeast rDNA transcription unit with marked processing sites.

### Figure S2

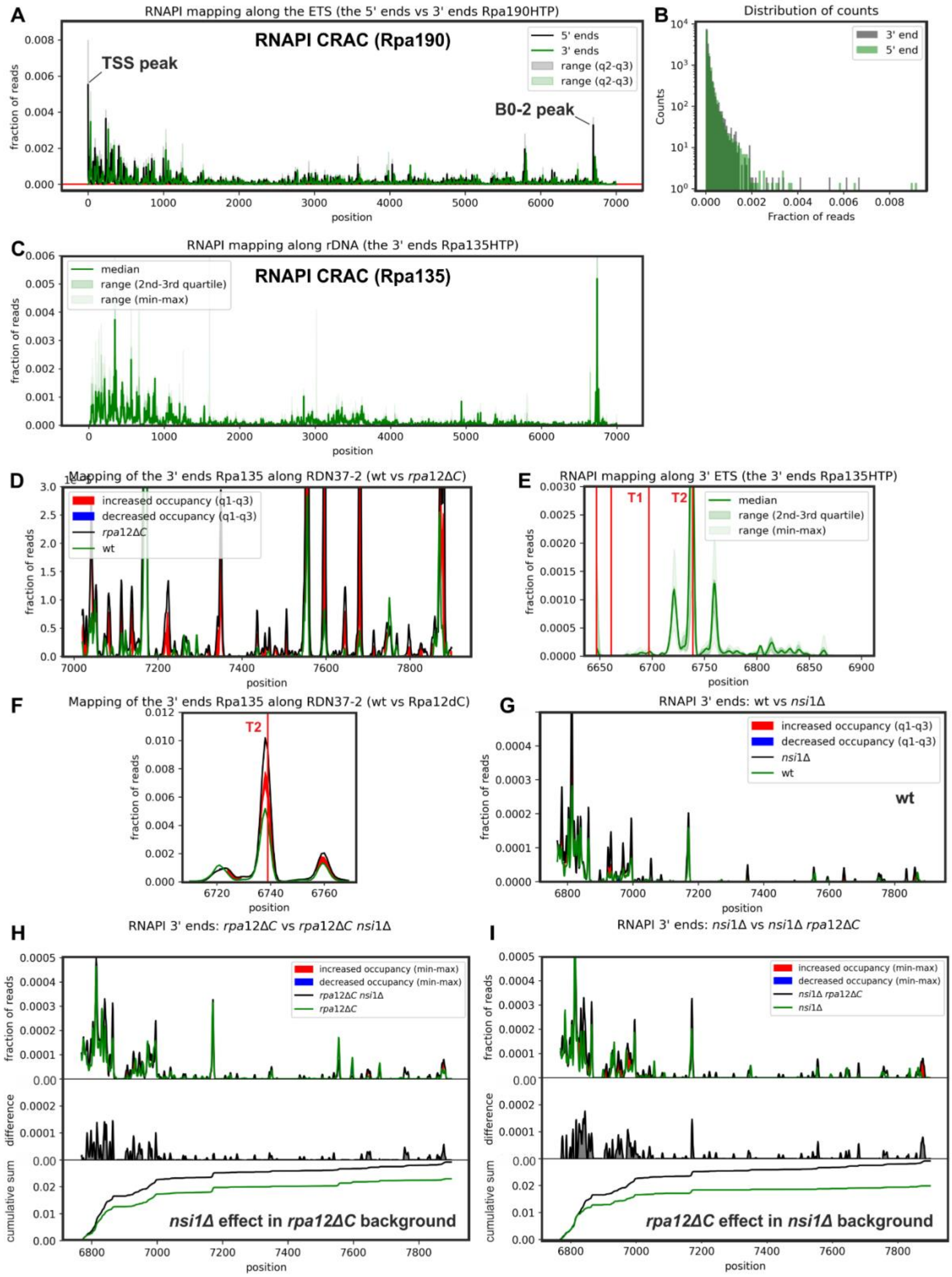


**Figure S2.** RNAPI termination is associated with decreased elongation kinetics.

A: RNAPI Rpa190-HTP CRAC mapping of the 5’ ends (black) and the 3’ ends (green) of reads. Note that substantial differences between the 5’ and 3’ ends are visible for the very 5’ end at the transcription start site (TSS peak) and within the 3’ ETS, at the B0-2 cleavage site (B0-2 peak).

B: 5’ ends of RNAPI CRAC reads are less distributed relative to 3’ ends.

C: Distribution of RNAPI Rpa135-HTP CRAC 3’ ends of reads along the rDNA.

D: Transcriptional read-trough is increased in *rpa12ΔC* strain in comparison to wt.

E: Distribution of RNAPI Rpa135-HTP CRAC reads around the 3’ end of the 35S pre-rRNA.

F: RNAPI Rpa135-HTP CRAC signal is increased at the T2 site in Rpa12ΔC strain.

G: Limited effect of *nsi1Δ* on RNAPI transcriptional read-trough detected with Rpa135-HTP CRAC in wt background.

H: Limited effect of *nsi1Δ* on RNAPI transcriptional read-trough detected with Rpa135-HTP CRAC in *rpa12ΔC* background.

I: Effect of *rpa12ΔC* on RNAPI transcriptional read-trough detected with Rpa135-HTP CRAC in *nsi1Δ* background.

### Figure S3

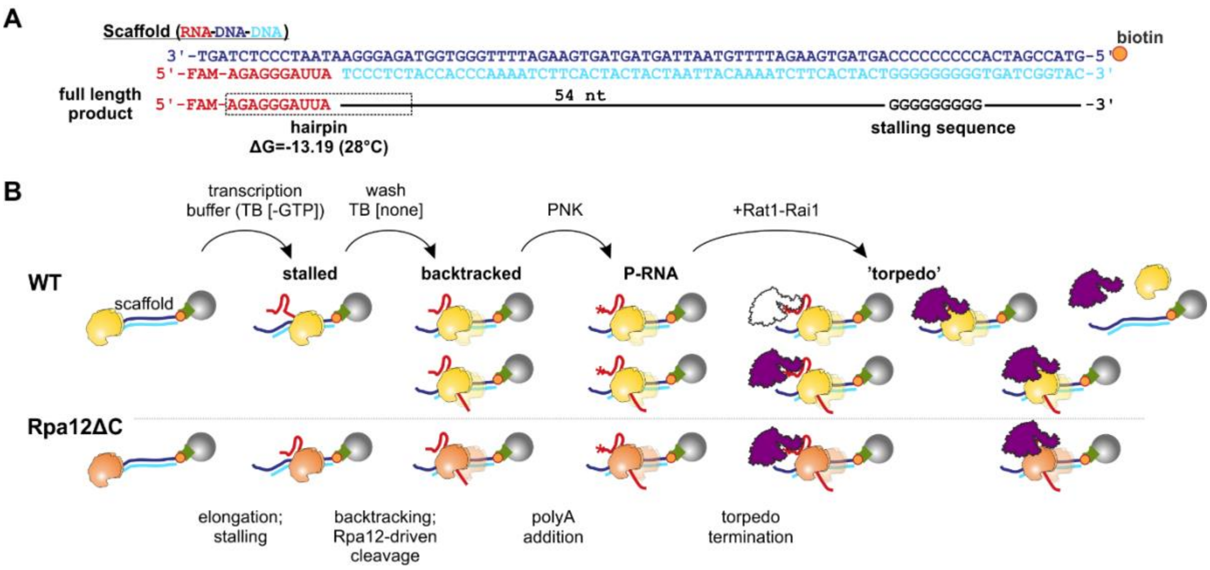


**Figure S3.** Schematic of *in vitro* termination assay.

A: Sequence of RNA-DNA-DNA scaffold immobilized via biotin-streptavidin interaction.

B: Schematic of the experiment.

### Figure S4

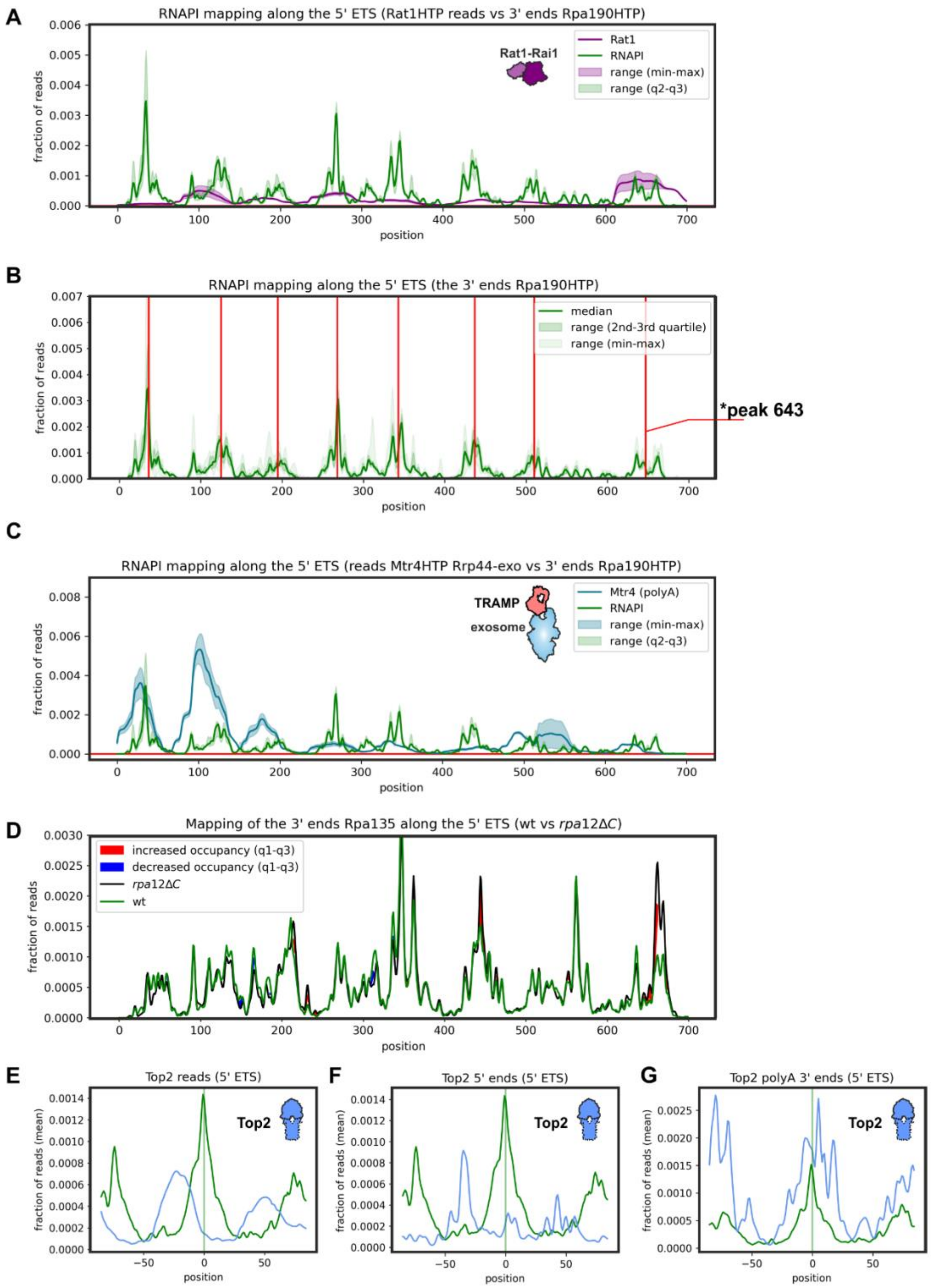


**Figure S4.** Rat1 and TRAMP are found at sites of slowed RNAPI elongation.

A: Rat1 CRAC reads superimposed with the 3’ ends of RNAPI CRAC reads.

B: Peaks found with peak-calling algorithm that were used to generate peak metaplot. * Note peak 643 was excluded from meta-analysis to separate Rat1 surveillance from canonical degradation of the 5’ ETS during pre-rRNA processing.

C: Mtr4 (Rrp44-exo) CRAC reads superimposed with the 3’ ends of RNAPI CRAC reads.

D: RNAPI (Rpa135-HTP) CRAC shows minor difference in *rpa12ΔC* strain in the 5’ ETS.

E-G: RNAPI Rpa190-HTP CRAC peak metaplot for the 5’ ETS, comparing Rpa190 and Top1-HTP CRAC at the level of single peaks: reads (E), the 5’ ends of reads (F) and the 3’ ends of polyadenylated reads (G).

**Figure S5**

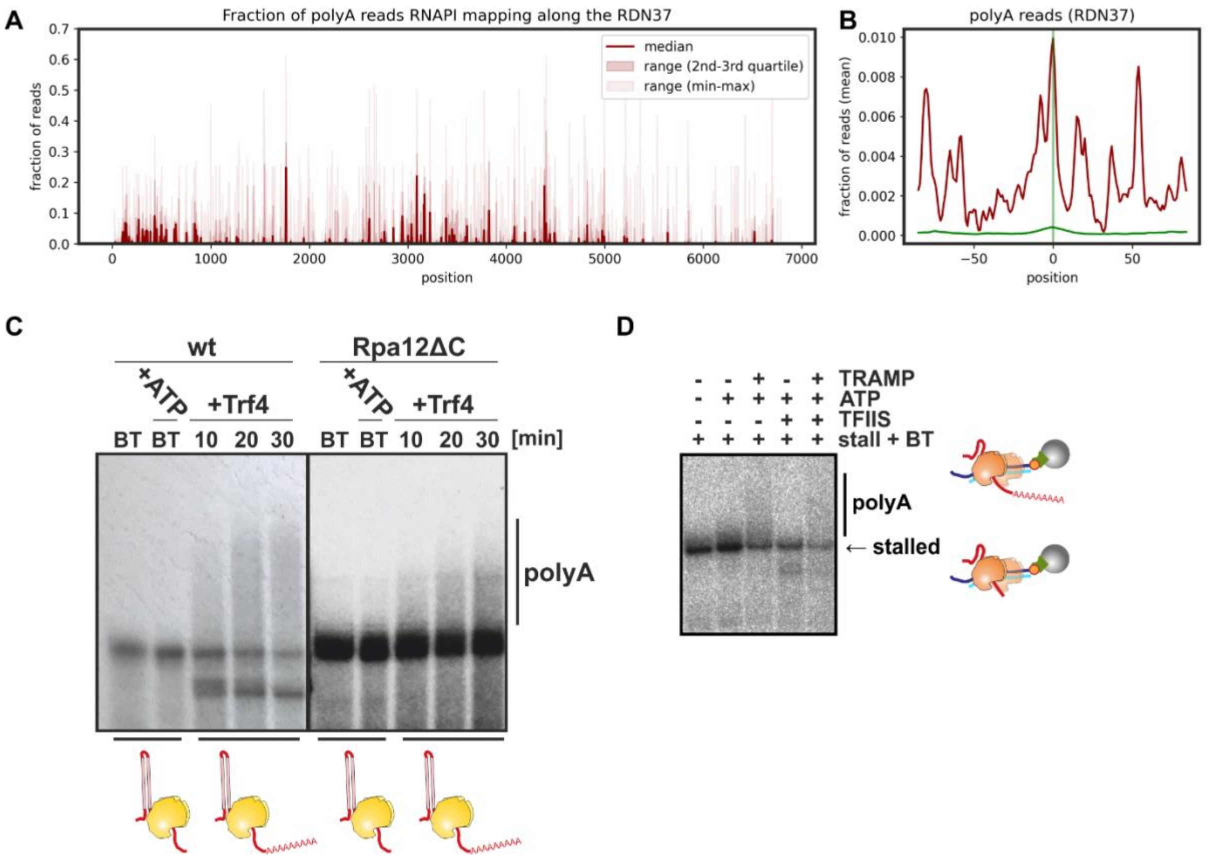


**Figure S5.** Backtracked nascent transcripts can be oligo-adenylated by the TRAMP complex

A: Fraction of polyA reads mapping along the *RDN37*.

B: RNAPI CRAC peak metaplot for *RDN37*, comparing the 3’ ends of the reads (green) with poly(A) reads.

C-D: Trf4 oligo-adenylates the 3’ end of backtracked, nascent RNA *in vitro* extruded from RNAPI (C) and RNAPII (D).

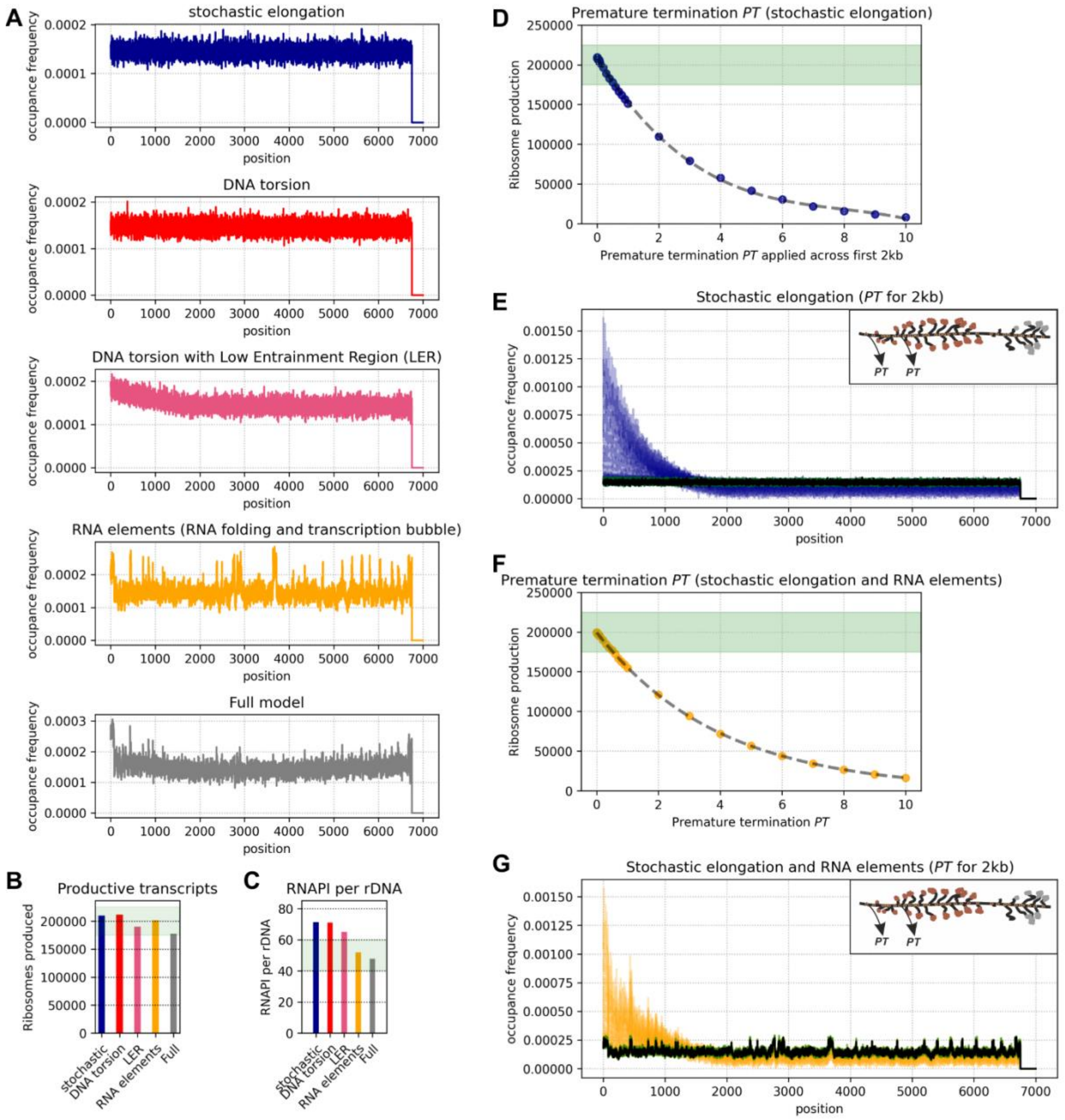


**Figure S6.** Development of premature termination $PT$ function.

A: Modeled RNAPI occupancy along the transcription unit using a model of stochastic initiation and discrete, stochastic elongation. The average RNAPI occupancy for 16 simulations is presented. Each simulation was run for 6,000 sec and 200 time points were collected.

B: Average number of productive transcription events for each type of simulation (corresponding to panel A).

C: The average number of RNAP molecules per transcription unit (corresponding to panel A).

D: $PT$ effect on ribosome production in model including stochastic elongation only. $PT$ was applied across the first 2kb of the transcription unit. Note that because $PT$ is applied only for this part of the transcription unit the values of $PT$ are significantly higher than for panel Fig 6A.

E: RNAPI occupancy profiles corresponding to panel D. Black – no premature termination $PT$, green $PT$ where ribosome synthesis is effective, blue – ribosome synthesis is decreased or insufficient.

F: $PT$ effect on ribosome production in model including stochastic elongation and RNA elements. Note that because $PT$ is applied only for the part of the transcription unit the values of $PT$ are significantly higher than for panel Fig. 6A.

G: RNAPI occupancy profiles corresponding to panel E. Black – no premature termination $PT$, green $PT$ where ribosome synthesis is effective, orange – ribosome synthesis is decreased or insufficient.

**Table S1.** Oligonucleotides used for strain construction.

| ID | Name | loc | sequence |
|---|---|---|---|
| PCR064 | Rpa190 check | fwd | TTTATCGACGTTGATGGTATTA |
| PCR065 | | rev | GCATTGACAGGCTCCATAA |
| PCR073 | Rpa135-HTP tagging | fwd | CTGAGCTATCCGCAATGGGTATAAGATTGCGTTATAATG<br>TAGAGCCCAAA<u>GAGCACCATCACCATCACC</u> |
| PCR074 | | rev | ACAATTGACCAAGCCTTCATTTACCATTCTATATCAATTTG<br>GAAAGAAGGGTA<u>TACGACTCACTATAGGG</u> |
| PCR075 | Rpa135 check | fwd | ATGTCGCGAGTGTGGTTCTATTTT |
| PCR076 | | rev | AAGCCTGCACCTCTTGCAGTAGAA |
| EP98 | Nsi1 F1 deletion | fwd | CAAATTTTGTGCATAGAGCAAGCAGCCGTTTCTTGTCT<br>GCGTCAAGAAGAAAGATAAAGGTAGACGGATCCCCGG<br>GTTAATTAA |
| EP99 | Nsi1 R1 deletion | rev | TTTAAAAATCAGTAAATATGCTTTTATCTATTGGGTCTGT<br>ATATGTTTGGGAAAGTAACCCTTCGAATTCGAGCTCGT<br>TTAAAC |
| EP100 | Nsi1 test | fwd | GAGCTTTCCAAATGCGATA |
| JH1584 | MX4-6 rev | | TGCAGCGAGGAGCCGTAAT |
| | Rai1 TAPfwd | fwd | TCTCGGAATAACAAGCAAAGTCGGTATGACAATTCAAG<br>AGCAAATAGGCGTTCCATGGAAAAGAGAAG |
| | Rai1Tap rev | rev | TTTATAAATTTGCGAAAACCTAAATTTACCATAAAATGC<br>GCACGAGTAGTTTACGACTCACTATAGGG |

**Table S2.** Oligonucleotides used for *in vitro* assay.

| ID | oligo | sequence |
|---|---|---|
| oTWT039 | 5'FAM | AGGCCGAAA |
| oTWT114 | | /56-FAM/rArGrArGrGrGrArUrUrA |
| oTWT150bio | | /5Biosg/ATACTTACAGCGTACCGATCACCCCCCCCAGTAGTGA<br>AGATTTTGTAATTAGTAGTAGTGAAGATTTTGGGTGGTAGAGGG<br>AATAATCCCTCTAGT |
| oTWT151 | | TCCCTCTACCACCCAAAATCTTCACTACTACTAATTACAAAATCT<br>TCACTACTGGGGGGGGTGATCGGTAC |
| oTWT165 | | rUrArUrArUrCrUrUrGrUrCrArArUrCrArUrArCrCrArGrArGrGrGrArUrUrA |
